# Systematic review and meta-analysis on the effects of chronic peri-adolescent cannabinoid exposure on schizophrenia-like behaviour in rodents

**DOI:** 10.1101/2023.12.15.571463

**Authors:** Zhikun Li, Diptendu Mukherjee, Bea Duric, Isabelle Austin-Zimmerman, Giulia Trotta, Edoardo Spinazzola, Diego Quattrone, Robin M Murray, Marta Di Forti

**Author notes:** **Corresponding author:** Dr Marta Di Forti, MD PhD, Social, Genetic and Developmental Psychiatry Centre, Institute of Psychiatry, Psychology and Neuroscience, King’s College London, London SE5 8AF, UK. These authors share first authorship (Zhikun Li, Diptendu Mukherjee).

## Abstract

**Background:** The link between cannabis use and schizophrenia is well-established in epidemiological studies, especially among adolescents with early-onset use. However, this association in rodent models is less clear. This meta-analysis examined the effects of adolescent cannabinoid exposure on distinct schizophrenia-like behaviours in rodents and how experimental variations influence outcomes.

**Methods:** Following a pre-registered protocol (CRD42022338761), we searched PubMed, Ovid Medline, Embse and APA PsychInfo for English-language original studies until 2022. We synthesised data from experiments on schizophrenia-like behaviour in rats and mice after repeated peri-pubertal (onset between P23-P45) cannabinoid exposure. Risk of bias was assessed using the SYRCLE’s tool.

**Results:** We included 291 experiments from 91 articles across 9 behavioural tests. We found meta-analytic evidence supporting that CB1R agonists, both natural and synthetic, elicited broad schizophrenia-like behavioural alterations, including impaired working memory (g =-0.58 [CI: -1.00,-0.16]), novel object recognition (g=-0.63 [CI: -0.97,-0.30]), novel object location recognition (g=-0.70 [CI: -1.22,-0.28]), social motivation (g=-0.40 [CI: -0.63, -0.16]), pre-pulse inhibition (g=-0.48 [CI: -0.89, -0.08]), and sucrose preference (g=-0.92 [CI: -1.87,0.04]). By contrast, effects on novelty-induced locomotion were negligible. Subgroup analyses revealed similar effects across sexes and species. Substantial variance in the protocols and moderate-to-high heterogeneity in behavioural outcomes were observed. We found CBD may attenuate novelty-induced locomotion in an open field and enhance fear memory recall, but data was limited.

**Discussion:** This is the first meta-analysis to comprehensively assess the link between cannabinoids and schizophrenia-like behaviours in rodents. Our results support epidemiological links between early cannabis use and schizophrenia-like phenotypes, confirming the utility of animal models. Standardising protocols will optimise models to strengthen reproducibility and comparisons, our work provides a framework for refining rodent models to elucidate biological pathways linking cannabis and schizophrenia.

## Introduction

Cannabis is one of the oldest and most widely used psychoactive substances in human history (1). The latest UN report estimated that the global number of cannabis users reached 209 million in 2020, representing a 23% increase from 2010 (2). The increasing popularity of cannabis use may be influenced by several factors, such as the legalisation of medicinal and recreational cannabis in some countries, the increased availability and social acceptance, and the perception that cannabis has low health risks (3,4). However, epidemiological studies have consistently shown a link between frequent cannabis use and psychosis (5,6). Recent evidence showed that daily cannabis users had a three-fold higher risk of developing psychosis than non-users. The risk increases with the use of high-potency cannabis that contains high levels of tetrahydrocannabinol (THC), the main psychoactive ingredient in cannabis (7). Furthermore, evidence suggested that earlier adolescent cannabis initiation may also confer greater psychosis vulnerability (8). Despite this knowledge, the biological mechanisms underlying this relationship, and the role of genetics have not been conclusively elucidated (9).

Human studies on cannabis and psychosis face considerable challenges in controlling for genetics, cannabis type, consumption patterns and social contexts. In contrast, rodent models enable controlled examination of cannabis exposure effects on the brain and behaviour. Given the conserved nature of brain circuits between humans and rodents (10,11) animal models hold promise for probing the pathophysiological mechanisms underlying cannabis effects relevant to humans.

Over the years, a growing body of research has investigated the impact of cannabinoids on schizophrenia-like behaviours in rodents (12–16). While providing useful preliminary evidence, these studies have substantial variability in factors like cannabinoid type and dosage, timing of exposure, sex, and species of animals used (14). Additionally, while numerous behavioural aspects have been reportedly to be impacted by cannabis exposure in rodents, the validity and sensitivity of the individual tests used to model complex schizophrenia-like behaviour in rodents remain unclear.

To address these questions, we conducted a systematic review and meta-analysis of rodent experiments that modelled chronic cannabis use and assessed its link with schizophrenia-like behaviours. We focused on studies that administered cannabinoids during adolescence, a critical neurodevelopmental period with heightened vulnerability to substance impacts (17–21). Through this meta-analysis, we aimed to 1) summarise existing behavioural data of rodent experiments that modelled adolescent cannabis exposure, 2) compare the impacts of distinct cannabinoids, particularly THC and cannabidiol (CBD), 3) explore the potential moderating factors of sex, species, age of treatment onset, time lapse between treatment and assessment (short-term vs long-term), and 4) discuss the implications for future research and identify open questions in the field of rodent models of cannabinoids exposure.

## Methods

This systematic review and meta-analysis followed the PRISMA (Preferred Reporting Items for Systematic Reviews and Meta-Analyses) guidelines (Supplementary appendix 1). The protocol was pre-registered at PROSPERO on 21^st^ June 2022, protocol number: CRD42022338761.

### Search Strategy

The literature search was conducted on 4^th^ June 2022 across electronic databases including PubMed, EMBASE, MEDLINE and APA PsycINFO, using the following keywords: 1) cannabis, 2) animal, and 3) adolescence. (Full search terms in Supplementary appendix 2). We included peer-reviewed original studies written in English.

### Study screening and eligibility criteria

The titles and abstracts of 2 341 articles retrieved from the preliminary search were screened and cross-checked by two independent reviewers (ZL and BD) and discrepancies were resolved through discussion with a third researcher (DM). Subsequently, the two reviewers evaluated the full texts of the remaining articles using specific inclusion criteria adapted from the PICO method (22) (Additional details in Supplementary appendix 3). We pre-specified 12 behavioural tests (Table 1) relevant to the three core domains of schizophrenia symptoms: positive symptoms, negative, and cognitive impairments (23–26). We pooled data for each test into separate meta-analyses.

**Table 1.** Behavioural tests and primary outcome measures for data extraction.

|  | Relevant schizophrenia symptoms |  |  |  |  |  |  |  |  |  |  |  |  |
| --- | --- | --- | --- | --- | --- | --- | --- | --- | --- | --- | --- | --- | --- |
|  | Positive symptoms |  | Social withdrawal |  | Working memory |  | Short-term memory |  | Sensori-motor gating | Associative memory | Spatial memory | Cognitive flexibility | Anhedonia |
| Test | Locomotion in open field |  | Social motivation | Social novelty preference | Y maze | T maze | Novel object recognition | Novel object location | Pre-pulse inhibition | Fear conditioning | Morris water maze | Attentional set-shifting | Sucrose preference |
|  | novelty induced | Psycho-stimulant-induced |  |  |  |  |  |  |  |  |  |  |  |
| Primary outcome measure | Total distance travelled in the open field |  | Social motivation index | Social preference index | %Correct alternations |  | Discrimination Index |  |  | %Time freezing in recall trial | Time to reach hidden platform | Number of trials to reach criterion in shift trial | Sucrose preference index |

### Data extraction strategy and quality assessment

Two reviewers (ZL and BD) independently extracted data using a standardised data extraction form (Supplementary Table 2). Importantly, we regarded a comparison between a control and a cannabinoid treatment group as a single experiment, and an effect size was calculated for each such comparison/experiment. Therefore, if an article included multiple control-treatment comparisons, each comparison was treated as one separate experiment, and multiple effect estimate data from that article were calculated and pooled.

Quality assessment was executed following the SYRCLE’s risk of bias tool for animal studies (27), which evaluated each article for their risk of bias across 10 items, assigning a rating of high risk, low risk, or unclear for each item.

### Statistical Analysis

Analyses were performed in R, using packages *meta*, *metafor* and *RevMan*. From each experiment, we calculated the effect sizes as Hedge’s *g*. The inverse variance-weighted random effects model with Knapp-Hartung adjustment was used to calculate overall effect sizes. The Restricted Maximum-Likelihood (REML) estimator was used for *τ*^2^.

The direction of effect sizes reflects the numerical change of effect in the experiment group compared to the control group. Data are presented as Hedge’s *g* ± 95% confidence intervals. Results were regarded as significant when the confidence interval entirely excluded zero and corresponded to a p value lower than 0.05 in Cochran’s Q test. A summary effect was considered valid only when at least 4 individual effect estimates could be pooled.

Heterogeneity was assessed using the *I^2^* statistics (28). We regarded an *I^2^* of 0% to 40%: might not be important; 30% to 60%: may represent moderate heterogeneity; 50% to 90%: may represent substantial heterogeneity; 75% to 100%: considerable heterogeneity (29).

The pre-specified list of subgroup analyses included assessments for the effect of a) species, b) sex, c) substance, d) time lapse of behavioural assessment post treatment. Cochran’s Q statistics was used to assess between-subgroup differences. We regarded a subgroup result as valid only when there was a minimum number of 4 experiments in each subgroup.

Publication bias was inspected qualitatively by assessing the asymmetry of funnel plots and quantitatively by Egger’s test (30). Sensitivity analyses were conducted to evaluate the robustness of the overall effect estimates when excluding studies a) with high-risk level of bias and b) reported alternative outcome measures.

## Results

A total of 2 341 records were identified through database search, amongst which 104 studies matched our inclusion criteria. 13 articles were subsequently excluded for insufficient data, resulting in n=91 studies published between 2003 and 2022 being used for quantitative data synthesis. [Figure 1A, C; Supplementary Data Sheet].

**Figure 1.**
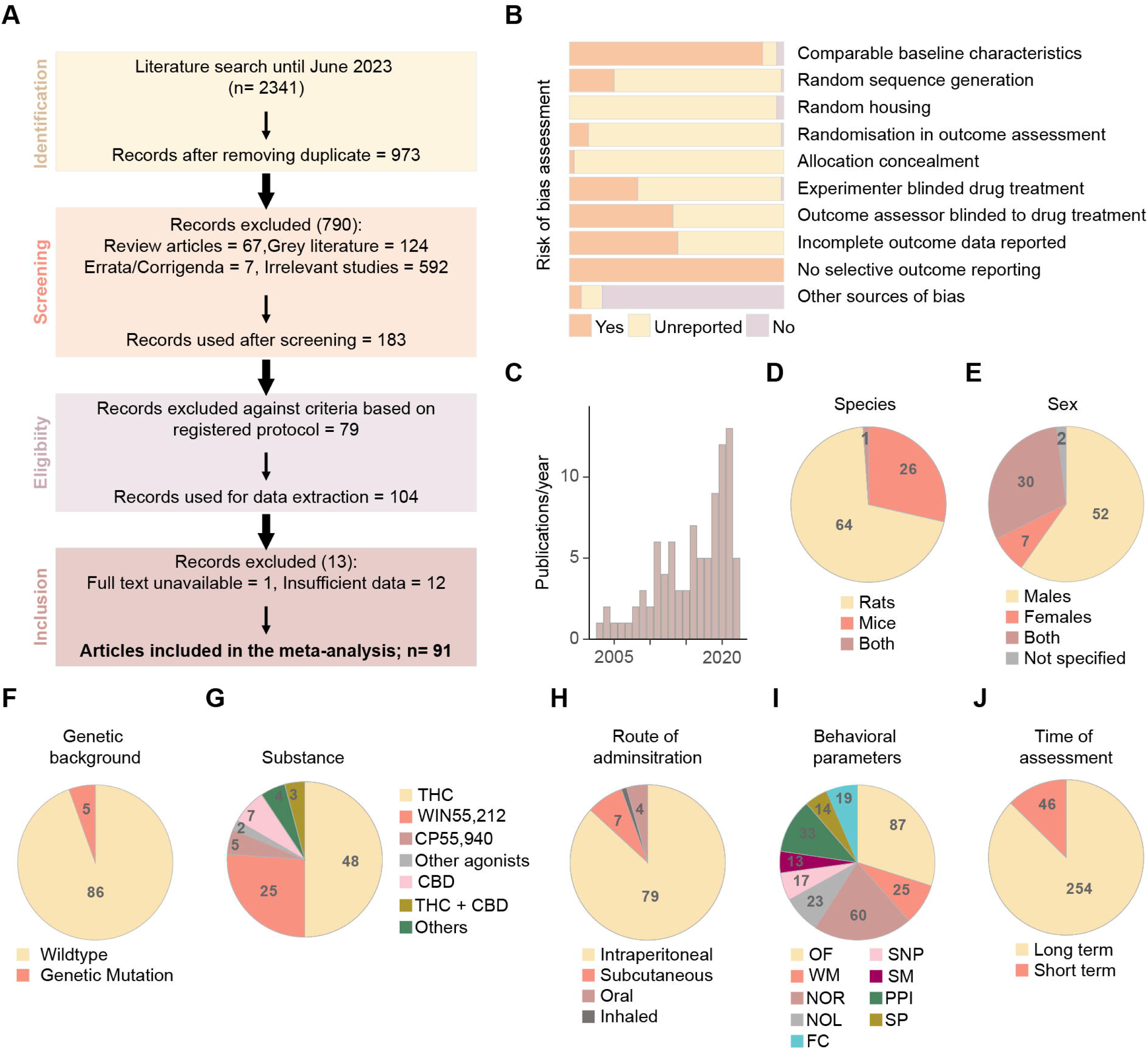
Article screening and general characteristics of the studies included. **A**. Flowchart of the screening strategy for the articles returned from electronic databases and the number of studies excluded at each step. **B**. Cumulative bar charts displaying risk of bias assessment across 10 items specified in SYRCLE’s assessment tool. **C**. Histogram showing the trend in articles published every year between 2003 and 2022 included in the meta-analysis. **D-J**: Pie charts depicting the proportion and number of studies categorized by species (D), sex (E), genetic background (F), substance administered (G), route of administration, and the proportion of experiments categorised by route of administration (H), behavioural parameters characterized (I) and time of behavioural assessment (short term: 24h to 10d after final dose; long term: > 10days after final dose) (J). OF- novelty-induced locomotion in an open field; WM- working memory; NOR- novel object recognition; NOL- novel object location; SNP- social novelty preference; PPI- prepulse inhibition, FC- fear conditioning; SM- social motivation; SP- sucrose preference.

Risk of bias assessment [Figure 1B] revealed all included articles pre-specified outcome measures and fully reported data. Most articles adequately controlled and reported baseline animal characteristics (90%) and provided sources of funding/conflict of interest statements (85%). However, few articles disclosed details on randomisation procedures and blinding methods as specified in the SYRCLE assessment tool. Therefore the majority of studies were labelled “unclear risks” for these domains.

To determine the diversity of experimental paradigms applied, we categorised studies according to the genetic background, species, sex, substance administered, route of drug administration, behavioural measures, and the timing of behavioural assessment. These data are presented as pie charts [Figure 1D-K]. We found that most studies (n=64) used rats as the model organism, while 26 studies used mice, and one study characterised both species [Figure 1D]. Over half of the studies included only one sex, with male animals (n=52) being preferred over females (n=7) [Figure 1E].

While the majority of the studies used wildtype animals, 5 studies used genetically modified mice [Figure 1F, a descriptive summary of gene-environment interaction findings in Supplementary appendix 4]. Regarding substances, the effects of THC (n=48) have been extensively profiled, followed by the synthetic full CB1R agonists WIN-55212-2 (n=25) and CP55,940 (n=5) [Fig 1G]. We also observed substantial variance in the tested THC doses (ranging from 0.2mg/kg to 15mg/kg), plus 75 experiments adopting an escalating paradigm (2.5-5-10mg/kg). Furthermore, non-contingent methods of drug delivery (31) like the intraperitoneal or subcutaneous injection (n=86) were more common than voluntary self-administration [Figure 1H].

Corresponding to the schizophrenia symptoms in humans, we specified a list of 12 behavioural tasks. From all the experiments (K=291) synthesised from the 91 articles, we registered 9 behavioural tests that were reported in more than 4 experiments. Morris water maze (K=3), attentional set-shifting (K=3), and psychostimulant-induced hyperactivity in an open field (K=3) were excluded due to insufficient outcome data. Amongst the 9 behavioural tests, novelty-induced locomotor activity in an open field test was most examined (K=87), followed by novel-object recognition (K=60) [Figure 1I]. Within our specified range of administration onset around puberty (P21-56), most studies began treatment in week 5 (P28-P34). Finally, the behavioural experiments were most frequently performed after an abstinent period of ten days (K=255), characterising primarily long-term effects of drug exposure [Figure 1J].

### Exposure to CB1R agonists and schizophrenia-like behaviours in rodents

Exposure to CB1R agonists was associated with a detrimental effect on working memory tasks [g =-0.58; 95% CI=(-1.00;-0.16), p=0.009; n=24] and short-term memory tests including novel object recognition [g=-0.63; 95%CI=(-0.97;-0.30), p <0.001, n=56], novel object location [g=-0.70; 95%CI=(-1.22;-0.28), p<0.01, n=22], social novelty preference [g=-0.80; 95%CI=(-1.32;-0.28), p <0.01, n=17], as well as sensorimotor gating assessed through pre-pulse inhibition [g=-0.48; 95%CI=(0.89;-0.08), p=0.02, n=27]. Rodents exposed to CB1R agonists also displayed behaviour related to negative symptoms, such as reduced social motivation [g=-0.40; 95%CI=(-0.63;-0.16), p < 0.005, n=12]. There was also a trend towards impaired 24h fear memory recall [g=-0.45; 95%CI=(-0.91; 0.00), p=0.051, n=11] and reduced sucrose preference [g=-0.92; 95%CI=(-1.87; 0.04), p=0.059, n=9]. Notably, novelty-induced locomotion in an open field was the only behaviour displaying no significant change following adolescent CB1R agonists treatment [g=-0.11; 95%CI=(-0.27; 0.05), p=0.17, n=75].

### Comparing behavioural effects of THC and Synthetic CB1R agonists

Synthetic Cannabinoids (SCs) encompass a broad class ofartificial compounds that mimic the effects of phytocannabinoids such as THC, but often at multiple times higher potency and binding affinity (32). In our meta-analysis, the SC agonists identified included WIN55,212-2, CP55,940, AB-PINACA, AB-FUBINACA, and 5-MDMB-PICA. In a subgroup analysis, we compared behavioural impacts of these SCs versus THC. Overall, THC and SCs produced similar effects [Fig.3]. The differences were non-significant for for novelty-induced locomotion in an open field (Q=0.02, p=0.88), working memory (Q=1.67, p=0.19), novel object recognition (Q=0.85, p=0.35), novel object location (Q=0.02, p=0.88), social novelty preference (Q= 0.01, p= 0.9), social motivation (Q= 2.91, p=0.08), and sucrose preference (Q= 0.01, p= 0.91). However, SCs showed greater impairment than THC on pre-pulse inhibition (Q= 7.51, p<0.05) and fear conditioning (Q= 18.92, p<0.0001).

### Exposure to CBD and behavioural modulations

In contrast to CB1R agonists, we found few studies assessing CBD’s impact that matched our inclusion parameters [Fig 1G, 2B]. Therefore, we analysed the effect sizes for a subset of behavioural parameters with at least 4 effect estimates pooled. [Figure 2A]. A small but significant negative effect was observed in novelty-induced locomotion in the open field test, indicating reduced locomotion from CBD pre-treatment [g=-0.23; 95%CI=(-0.44; -0.01), p<0.05, n=12]. We also observed significant effects of CBD in enhancing fear memory retrieval [g=0.53; 95%CI= (0.04;1.02), p<0.05, n=8]. No significant effect was observed for novel object recognition [g=-0.44, 95%CI= (-1.64; 0.76), p=0.32, n=4] pre-pulse inhibition [g=0.40, 95%CI=(-0.60; 1.41), p=0.34, n=6] and sucrose preference tests [g=0.10, 95%CI=(-0.75; 0.95), p=0.77, n=5] with CBD.

**Figure 2.**
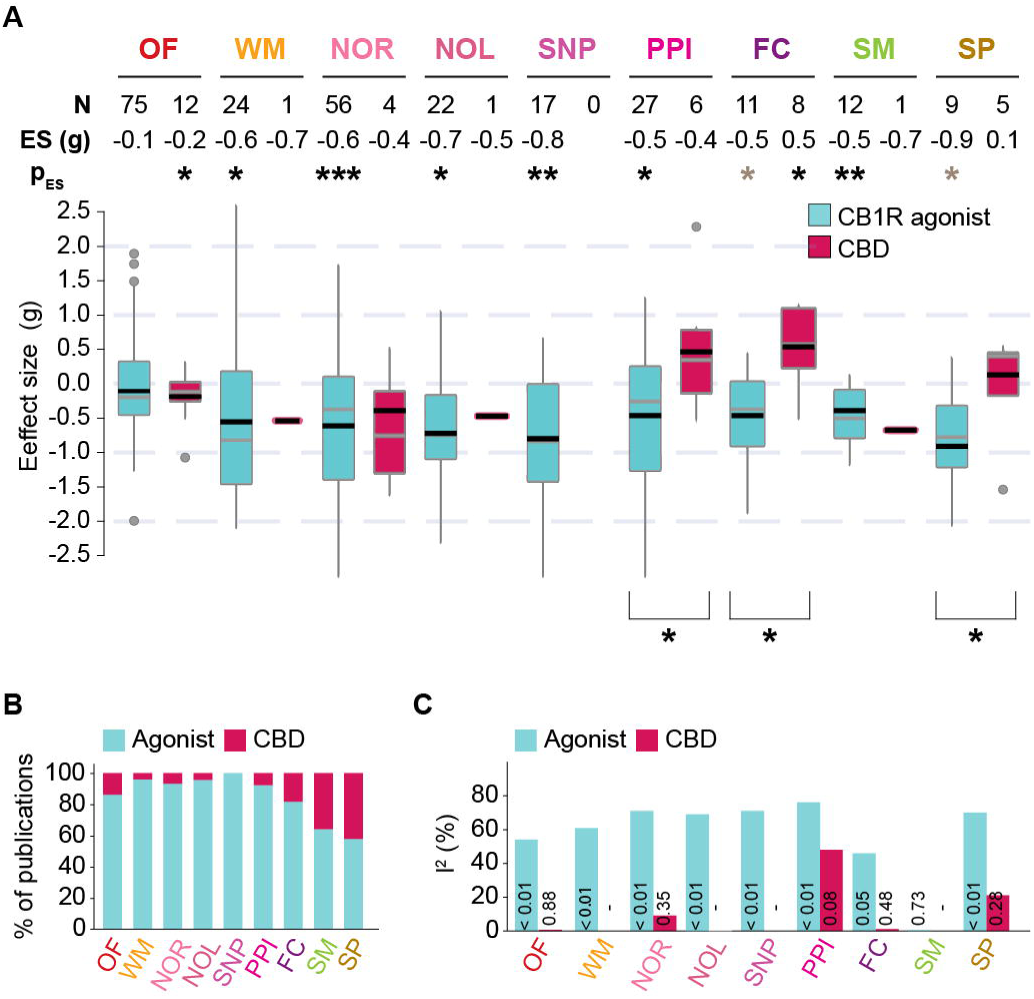
Chronic adolescent exposure to CB1R agonists and CBD differently modulates schizophrenia-like behaviour. **A**. Boxplots displaying the distribution of effect sizes (Hedge’s *g*) of CB1R agonists (blue) and CBD (red) across nine behavioural tests; Number of experiments included (N), values of pooled effect sizes (ES) and corresponding significance levels (P_E_S) are listed on top of each corresponding boxplot. Cochran’s Q statistics to compare effect sizes between drug interventions are denoted by asterisks below the boxplots. * p<0.05, **p<0.005, *** p<0.0005. Black horizontal lines in the boxplots indicate pooled effect sizes (inverse variance weighted, random effect model), grey horizontal lines indicate medians. Outliers are indicated by grey dots. **B**. percentage of publications analysing CB1R agonists (blue) and CBD (red) in each behavioural test. **C**. Bar charts displaying heterogeneity of each analysis represented by *I^2^* statistic and associated p values shown within each bar. OF-locomotion in a novel open field; WM-working memory; NOR-novel object recognition; NOL- novel object location; SNP- social novelty preference; PPI- prepulse inhibition, FC- fear conditioning; SM- social motivation; SP- sucrose preference. Forest plots for each behavioural test are available in supplementary figures S3-15.

**Figure 3.**
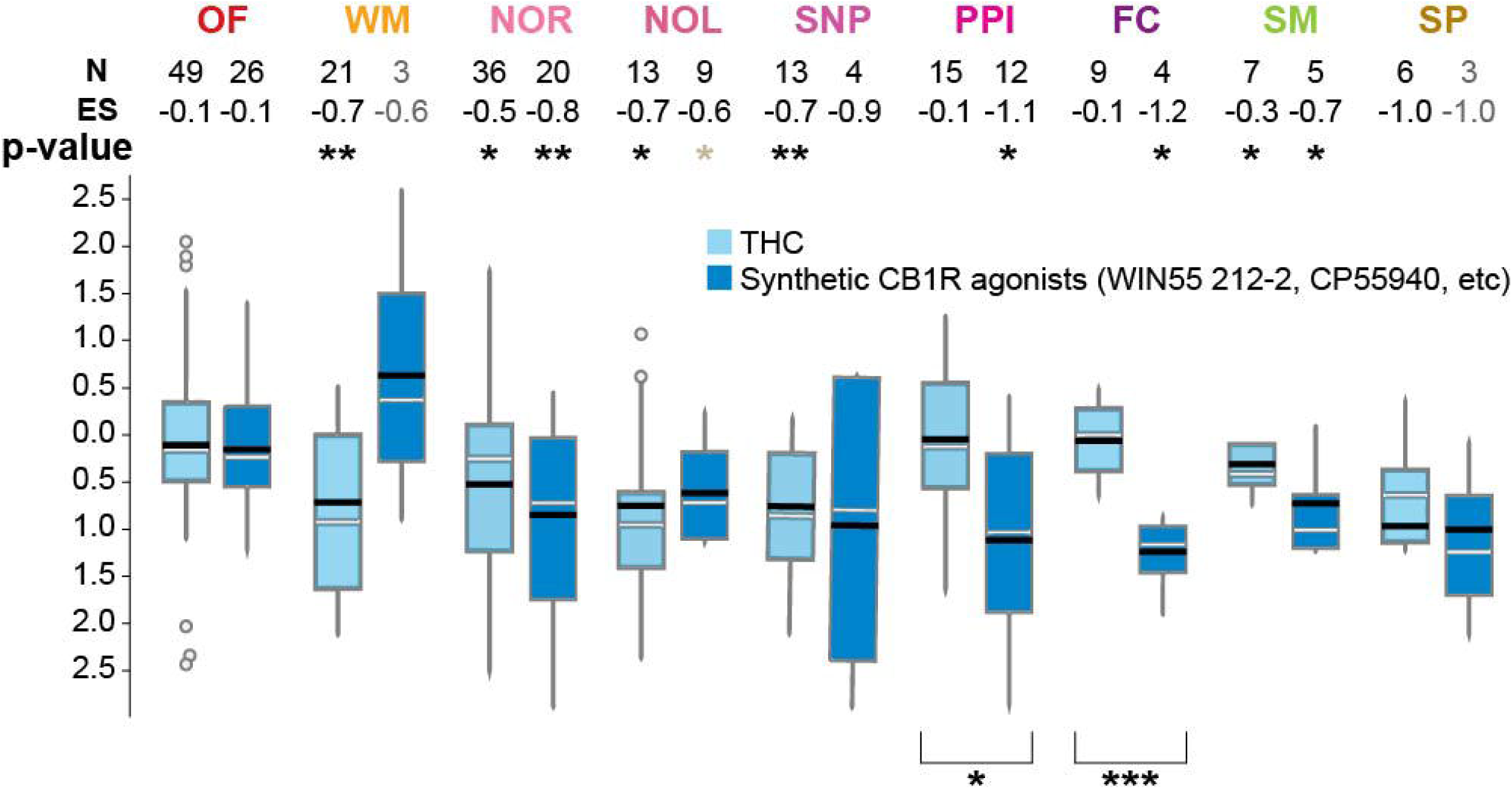
THC and synthetic CB1R agonists are associated with similar schizophrenia-like behavioural modification. A. Boxplots displaying the distribution of effect estimates (Hedge’s *g*) of THC (light blue) and synthetic CB1R agonists (dark blue) for different behavioural tests; Number of experiments included (N), pooled effect size (ES) and corresponding significance levels are listed on top of each corresponding boxplot. Significant subgroup differences between THC and SC estimated by Cochran’s Q statistics are denoted by asterisks below the boxplots. * p<0.05, **p<0.005, *** p<0.0005. Black horizontal lines in the boxplots indicate pooled effect sizes (inverse variance weighted, mixed effect model), grey horizontal lines indicate medians. Outliers are indicated by grey circles. OF-locomotion in a novel open field; WM-working memory; NOR-novel object recognition; NOL- novel object location; SNP- social novelty preference; PPI- prepulse inhibition, FC- fear conditioning; SM- social motivation; SP- sucrose preference.

Subsequently, comparing the effects of CB1R agonists and CBD, we found a significant difference for pre-pulse inhibition (Q=4.09, p<0.05), sucrose preference (Q=3.85, p<0.05) and fear memory retrieval (Q=11.32, p<0.005). In each of these tests, CBD improved the performance, whereas CB1R agonists worsened it. No difference was detected in the novel object recognition test (Q= 0.23, p=0.63) or novelty-induced locomotion in an open field (Q=1.06, p=0.30).

### Heterogeneity, publication bias and sensitivity analysis

We identified moderate to high levels of heterogeneity for most behavioural tests, with *I^2^* statistics ranging between 46% and 76%, except for the social motivation test [Figure 2C]. To identify potential sources, statistical outlier analyses were conducted. Outliers were defined as studies in which the 95% confidence interval of the effect size did not overlap with the confidence interval of the pooled effect. Statistical outliers were detected in most behavioural tests, accounting for a portion of the heterogeneity identified for each of the tests (Supplementary appendix 5).

We found significant asymmetry in 5 out of 14 funnel plots, indicating publication bias (4 out of 9 behavioural outcomes measured for CB1R agonist and 1 out of 5 for CBD) [Figure 4]. These include novel object recognition [intercept=-3.683, 95%CI=(-5.11; -2.25), p<0.001], novel object location [intercept=-5.495; 95%CI=(-8.96; -2.03), p<0.05], social motivation tests [intercept=-2.125, 95%CI=(-3.67;-0.58), p<0.05] for CB1R agonists, and sucrose preference test for CB1R agonists [intercept= 6.265, 95%CI=(-8.54; -3.99), p<0.05] and pre-pulse inhibition for CBD [intercept=15.371, 95%CI=(6.44; 24.31), p<0.05], indicating the existence of publication bias.

**Figure 4.**
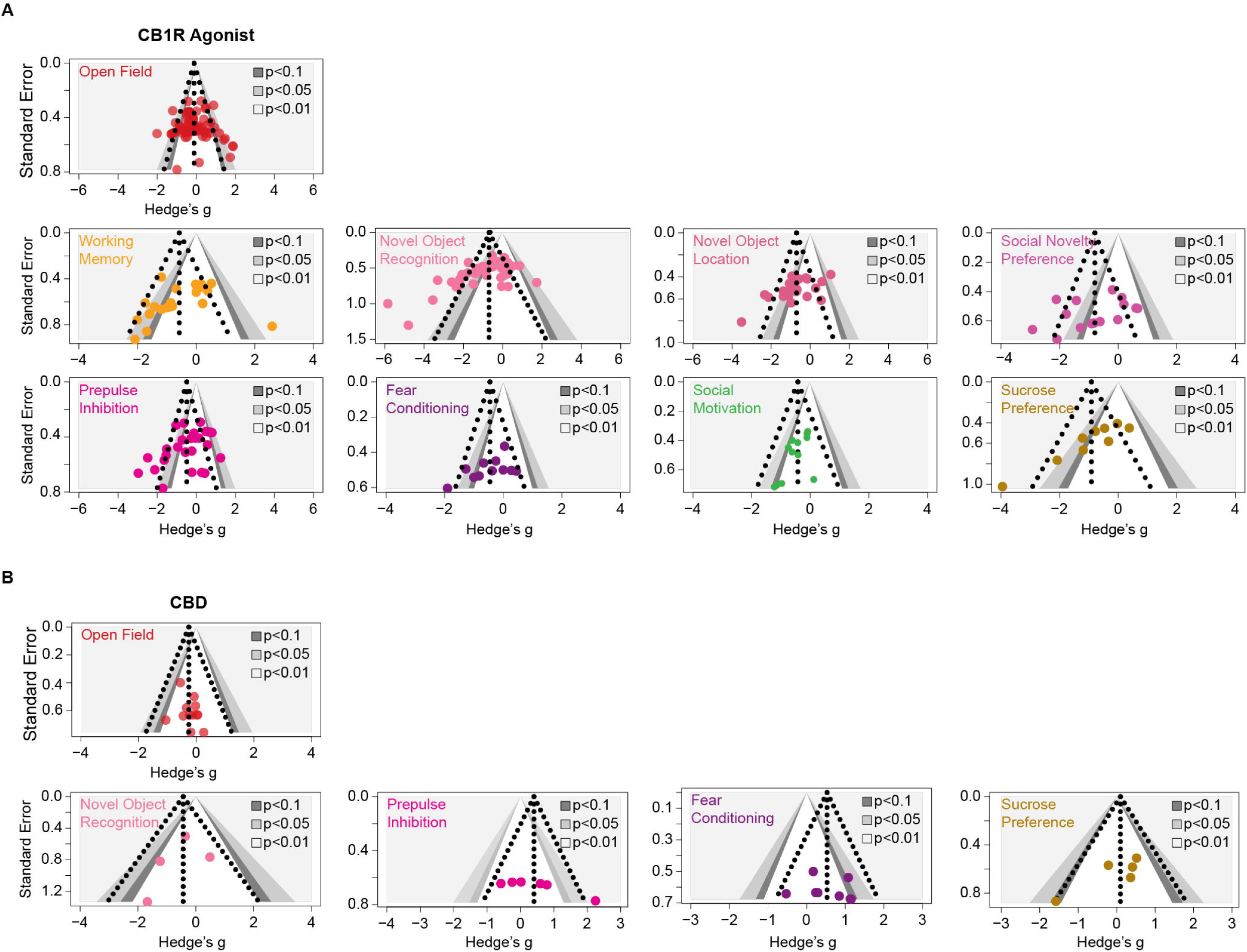
Contour-enhanced funnel plot for assessing potential publication bias for each individual outcome. Effect sizes (Hedge’s *g*) are plotted on the x-axis and standard errors on an inverted y-axis. Each point on the plot represents a single effect size data point. The dashed funnel is the idealised shape that we expect our studies to follow. The vertical line in the middle of the funnel shows the average effect size. The contours from light to dark grey signify the significance levels of the data points.

Furthermore, we performed a sensitivity analysis to assess whether the inclusion of alternative outcome measures would significantly modify the overall effects. Among the 9 behavioural tests, only novelty-induced locomotion in an open field included alternative outcome measures (e.g., beam breaks, number of floor squares entered) in addition to the primary measure (total distance travelled) for locomotor activity. The analysis result showed no significant change in pooled effect size before and after removing alternative measures. (ES_total_= -0.12, ES _primary_= -0.21, Q=0.46, p=0.50). However, sensitivity analysis based on risk of bias was not feasible, as the SYRCLE tool does not generate an overall high or low bias rating, and most included studies lacked sufficient information for risk judgement.

### Behavioural outcomes of chronic adolescent CB1R agonist exposure across rodent species and sexes

To further explore how variations in experiment design might impact behavioural outcomes and contribute to between-study heterogeneities, we performed a series of subgroup analyses. All subgroup analyses were performed on data obtained from studies on exposure to CB1R agonists, as CBD studies were sparse.

We first compared the subgroup effect sizes for mice and rats [Figure 5A]. Significant between-species difference was found only in fear conditioning recall (Q=7.25, p<0.05), albeit all experiments in the rat subgroup came from the same article, which bore the risk of being skewed by the same unidentified confounds. Heterogeneity remained moderate to high in the subgroups, indicating that species was not a significant moderator in our meta-analysis.

**Figure 5.**
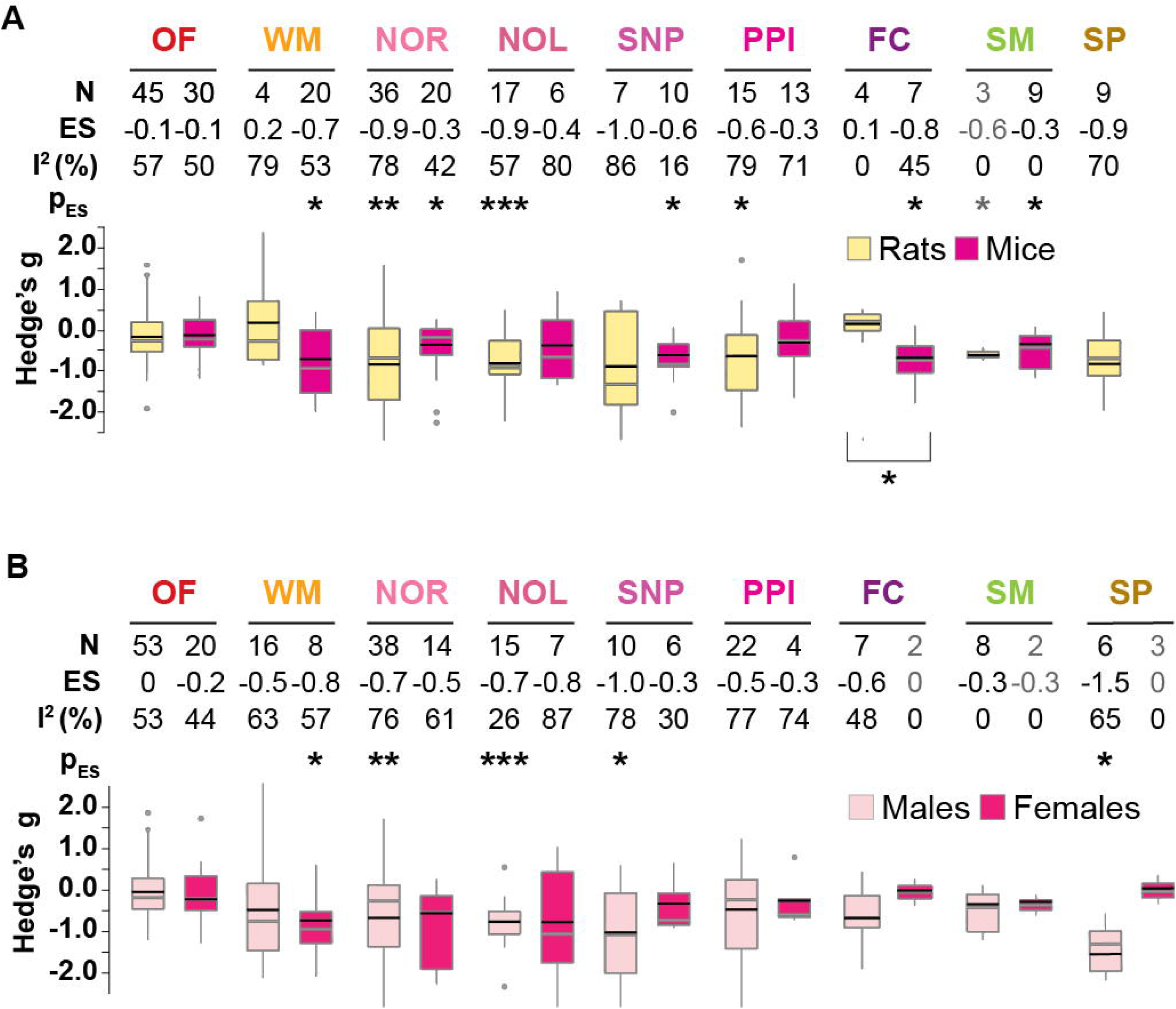
Boxplots displaying the distribution of effect sizes (Hedge’s g) for CB1R agonists in each behavioural test sub-grouped by **A.** species and **B.** sex. Number of experiments (N), pooled effect size values (ES), corresponding *I^2^*statistics and P values are shown on the top of each corresponding boxplot. Black horizontal lines in the boxplots indicate pooled subgroup effect sizes (inverse variance weighted, mixed effect model), grey horizontal lines indicate medians. Outliers are indicated by grey dots. Significant between-subgroup differences are denoted by asterisks below the boxplots. Comparing the effect sizes between rats and mice: OF-locomotion in a novel open field: Q=0.08,p=0.77; WM- Working memory: Q=1.32, p=0.25; NOR- Novel object recognition: Q=1.82, p=0.18; NOL- Novel object location: Q=1.18, p=0.28; SNP- Social novelty preference: Q=0.16, p=0.69; PPI- Prepulse inhibition: Q=0.87, p=0.35; FC- Fear conditioning: Q=7.25, p<0.05; SM- Social motivation, SP- sucrose preference. Comparing the effect sizes between males and females: OF- Q= 0.49; p=0.49, WM- Q=0.68; p=0.4, NOR- Q=0.21; p=0.64, NOL- Q=0.68; p=0.4, SNP- social novelty preference- Q=1.85; p=0.17, PPI- Q=0.20; p=0.65. * p<0.05, **p<0.005, *** p<0.0005.

Figure 5B displays the subgroup effect sizes for each behavioural test by sex. Notably, none of the behavioural tests revealed significant differences between subgroups. Furthermore, only the female subgroup for social novelty preference showed a substantial reduction in heterogeneity (*I^2^*=30%) relative to the overall effect.

### Comparing short-term and long-term effects of CB1R agonist exposure

To distinguish the short-term and protracted effects of CB1R agonists, we stratified the data based on whether the behavioural tests were performed between 24 hours to 10 days after the final dose or after an abstinence period of more than 10 days. As shown in Fig. 1K, most studies assessed behaviour exclusively in the long term. Therefore, we conducted subgroup analyses only for novelty-induced locomotion and novel object recognition. Substantial between-subgroup difference was found for novelty-induced locomotion (Q=10.09, p<0.005), where CB1R agonists administration was linked with significantly reduced locomotion in the short-term [g= -0.66, 95%CI= (-0.96; -0.36), n= 14, p<0.005, *I^2^*= 15%] but not in the long-term [g= -0.77, 95% CI = (-0.24; 0.10), n=68, p= 0.43, *I^2^* = 54%]; By contrast, performance on novel object recognition was shown significantly impaired in the long-term [g = -0.66, 95% CI = (-1.03; -0.29), n= 52, p<0.005, *I^2^*= 73%] but not evident in the short-term [g= -0.71, 95% CI = (-1.66; 0.24), n= 6, p=0.11, *I^2^*=67%]. However, heterogeneity remained high in the subgroups, and the subgroup difference was not statistically significant for novel object recognition (Q= 0.01, p= 0.90).

## Discussion

Previous meta-analyses have synthesised preclinical evidence for cannabinoids effects on rodent behaviour related to nociception (33), sleep (34) anxiety and depression (35,36), and some narrative systematic reviews have addressed aspects of schizophrenia-related behaviour such as social behaviour (37) and cognitive function (38). However, to our knowledge, this is the first systematic review to meta-analyse results from a comprehensive battery of tests for schizophrenia-like behaviour, focusing on adolescent cannabinoids exposure.

Our meta-analysis revealed a robust association between adolescent exposure to natural and synthetic CB1R agonists and impaired schizophrenia-related behavioural phenotypes in rodents. We report that exposure to CB1R agonists is associated with prominent cognitive deficits and pronounced behaviour changes similar to positive and negative symptoms of schizophrenia. Notably, these effects were persistent even after long-term abstinence. This suggests that adolescent exposure to CB1R agonists may cause lasting disruptions to the brain and impaired behaviour that extends into adulthood in rodents. Our results are consistent with the conclusion from existing reviews on rodent literature (15,39,40).

### Locomotor hyperactivity as a proxy for positive symptoms of schizophrenia

Novelty-induced hyperactivity in an open field is often considered a proxy for positive symptoms of schizophrenia (41,42) due to the well-established connection between dopamine and movement control(43–46). Indeed, enhanced striatal and subcortical dopaminergic activity has been reported in schizophrenia patients (47–50) and rodents (51,52). Although dopaminergic dysfunction is not the only pathophysiological mechanism affected, it is proposed to be a final common pathway where multiple effector pathways converge (53,54) Therefore, modelling a behaviour that is susceptible to dopamine-dependent changes provides construct validity to this paradigm.

However, we found that adolescent exposure to CB1R agonists did not have a significant overall effect on novelty-induced locomotor activity, and this was not modified by sex or species. In a subgroup analysis separating short-term versus long-term effects, we found a significant locomotor suppressing effect when the test was performed more recently to treatment cessation. This could be due to some residual effects of the acute drug-induced hypoactivity commonly observed immediately after drug administration, which subsided during abstinence.

However, the lack of evidence of a long-term effect on locomotion does not necessarily negate the hypothesis that CB1R activation modifies dopaminergic signalling(55–57). In fact, chronic long-term exposure to CB1R agonists could have a paradoxical effect on dopamine. Similar to other drug dependence models such as with psychostimulants or opioids (58,59), it has been shown that chronic cannabis users often display suppressed dopamine release (60–62). Furthermore, striatal dopamine levels are also found to be reduced in patients with dual diagnoses, including schizophrenia patients with a history of cannabis use under controlled and stressed conditions (63,64). These observations would suggest that using a simple locomotion test to model positive symptoms in cannabis-treated rodents oversimplifies a complex relationship. Therefore, we suggest refined protocols to study positive symptoms are necessary. Over the years, several such protocols have been developed, such as models of “altered reality testing” through Pavlovian conditioning (65,66), artificial manipulation of perceptual decision making (67,68), and a more recent development which probes experimentally controlled auditory hallucinations in rodents (69).

### CBD in schizophrenia

In this meta-analysis, we identified 7 studies that examined the effects of CBD on schizophrenia-like behaviour in wildtype animals. We found that chronic adolescent treatment of CBD was associated with a moderate but significant effect on reducing novelty-induced locomotion in the open field, and on improving fear memory recall in the fear conditioning task. However, results from other tests were non-significant and likely underpowered.

While the effects of CBD alone might be less prominent, some preclinical studies show that CBD reduced hyperlocomotion induced by psychotomimetic agents like amphetamine and ketamine (70), as well as in glutamatergic dysfunction models of schizophrenia (71). Social and sensorimotor deficits were also shown to be alleviated by chronic CBD treatment in MK801-induced schizophrenia models (71) and in stress-induced models (72,73). Along the same lines, it has been proposed that CBD may mitigate the psychotic effects of CB1R agonists by acting as a negative allosteric modulator of CB1R activity (74,75), and/or modifying downstream signalling (76). Here, we identified a need for more research to elucidate the impact of CBD and THC co-administration during adolescence, as existing evidence is limited and inconsistent.

### Sex-mediated effect of cannabinoids in rodents

Preclinical and clinical studies have consistently reported sex-dimorphic effects of cannabinoids (77,78). However, subgroup analyses from the current meta-analysis did not reveal sex as a significant mediator. This result should be interpreted with caution, as the uneven data distribution and low statistical power of our subgroup analysis could explain the lack of sex-mediated effect. We also noticed that although sexually dependent effects were frequently observed among individual studies, the direction and magnitude often varied greatly, sometimes contradictory. Therefore, an average effect of these studies might not be informative. Herein, we echo the calls for more inclusion of female animals in preclinical research (79–81), as this is essential for improving its clinical translation.

### Limitations and challenges

This meta-analysis provides a rigorous and comprehensive assessment of the effects of adolescent cannabinoid exposure on rodent behaviour. However, we must consider the limitations of our study. Firstly, the great disparity in the experimental settings of the animal studies, such as the age of exposure and the treatment protocols, could profoundly influence the animals’ response to drugs, which in turn increases heterogeneity and reduces the validity and generalizability of our findings. To circumvent this, we had initially planned additional subgroup analyses to explore the impact of dose and age of onset. However, such an effort was restricted due to the lack of sufficient data, and no conclusive association could be observed. For reference data, see supplementary Figure S1 and S2.

Second, many studies did not report essential information on experimental design and outcome data adequately. This hindered the risk of bias assessment and sensitivity analysis, and the outcome data had to be frequently extracted using a digital ruler software from graphical representations. Third, we could not include three behavioural tests for schizophrenia-like behaviour in the quantitative synthesis, because they either had insufficient data reported, or they lacked standardised protocol and outcome measurement. Fourth, some of our subgroups included data from multiple experiments within the same study rather than from different studies. The potential lack of independence could impact the accuracy of the overall effect estimates due to uncontrolled study-specific factors. Finally, the publication bias observed in our meta-analyses may indicate selective reporting of significant data, which could potentially result in inflated or biased summary effects.

### Future directions

In conducting this meta-analysis, we found that the definition of adolescence in rodents varied greatly, with a window ranging from P23 to P45. Adolescence marks a critical period for neurodevelopmental changes, when the brain structure and function change immensely, and are highly susceptible to the effects of drugs (40,82,83). Therefore, standardising the adolescent period would improve comparability and interpretability. Another key challenge in studying the effects of cannabis on schizophrenia-like behaviour is to devise a drug delivery method that accurately reflects human cannabis use. Most of the current studies rely on the intraperitoneal route, which is convenient but may not capture the complexity and variability of human cannabis consumption patterns. We are encouraged by recent studies utilising more translationally-relevant models like inhaled cannabis vapour (84), but such studies remain uncommon. Moving forward, characterising cross-species pharmacodynamic/pharmacokinetic correlations for cannabinoids would enhance model validity and reliability. Investigating dose-response relationships and drug interactions would also inform model optimisation. Such efforts to improve translational relevance will accelerate insights into mechanisms linking human cannabis exposure and schizophrenia phenotypes, informing prevention and treatment.

In conclusion, despite substantial variation in experimental protocols and paradigms, the results of this meta-analysis confirm that chronic exposure to CB1R agonists during adolescence is associated with the expression of some schizophrenia-like behaviours in rodents. This supports findings from human epidemiological studies. Moving forward, standardization of protocols, consideration of developmental periods and sex differences, and inclusion of diverse cannabinoid combinations will facilitate cross-species translation to elucidate the mechanisms linking adolescent cannabis use and schizophrenia phenotypes.

## Data availability

The data supporting the findings of this study are available within this article and the Supplementary Data Sheet.

