## Supplementary materials for "Systematic review and meta-analysis on the effects of chronic peri-adolescent cannabinoid exposure on schizophrenia-like behaviour in rodents"

**Supplementary appendix 1. PRISMA checklist**

| **Section and Topic** | **Item #** | **Checklist item** | **Location where item is reported** |
| --- | --- | --- | --- |
| **TITLE** | | |  |
| Title | 1 | Identify the report as a systematic review. | 1 |
| **ABSTRACT** | | |  |
| Abstract | 2 | See the PRISMA 2020 for Abstracts checklist. | 2 |
| **INTRODUCTION** | | |  |
| Rationale | 3 | Describe the rationale for the review in the context of existing knowledge. | 3-4 |
| Objectives | 4 | Provide an explicit statement of the objective(s) or question(s) the review addresses. | 3-4 |
| **METHODS** | | |  |
| Eligibility criteria | 5 | Specify the inclusion and exclusion criteria for the review and how studies were grouped for the syntheses. | 4; Supplementary appendix 3 |
| Information sources | 6 | Specify all databases, registers, websites, organisations, reference lists and other sources searched or consulted to identify studies. Specify the date when each source was last searched or consulted. | 4; Supplementary appendix 2 |
| Search strategy | 7 | Present the full search strategies for all databases, registers and websites, including any filters and limits used. | 4; Supplementary appendix 2 |
| Selection process | 8 | Specify the methods used to decide whether a study met the inclusion criteria of the review, including how many reviewers screened each record and each report retrieved, whether they worked independently, and if applicable, details of automation tools used in the process. | 4; Supplementary appendix 2 |
| Data collection process | 9 | Specify the methods used to collect data from reports, including how many reviewers collected data from each report, whether they worked independently, any processes for obtaining or confirming data from study investigators, and if applicable, details of automation tools used in the process. | 4-5;  Supplementary appendix 2 |
| Data items | 10a | List and define all outcomes for which data were sought. Specify whether all results that were compatible with each outcome domain in each study were sought (e.g. for all measures, time points, analyses), and if not, the methods used to decide which results to collect. | 4; Table 1; Supplementary appendix 2; |
|  | 10b | List and define all other variables for which data were sought (e.g. participant and intervention characteristics, funding sources). Describe any assumptions made about any missing or unclear information. | Data Extraction Table |
| Study risk of bias assessment | 11 | Specify the methods used to assess risk of bias in the included studies, including details of the tool(s) used, how many reviewers assessed each study and whether they worked independently, and if applicable, details of automation tools used in the process. | 5 |
| Effect measures | 12 | Specify for each outcome the effect measure(s) (e.g. risk ratio, mean difference) used in the synthesis or presentation of results. | 5 |
| Synthesis methods | 13a | Describe the processes used to decide which studies were eligible for each synthesis (e.g. tabulating the study intervention characteristics and comparing against the planned groups for each synthesis (item #5)). | Supplementary appendix 2 |
|  | 13b | Describe any methods required to prepare the data for presentation or synthesis, such as handling of missing summary statistics, or data conversions. | Supplementary appendix 2 |
|  | 13c | Describe any methods used to tabulate or visually display results of individual studies and syntheses. |  |
|  | 13d | Describe any methods used to synthesize results and provide a rationale for the choice(s). If meta-analysis was performed, describe the model(s), method(s) to identify the presence and extent of statistical heterogeneity, and software package(s) used. | 5 |
|  | 13e | Describe any methods used to explore possible causes of heterogeneity among study results (e.g. subgroup analysis, meta-regression). | 5 |
|  | 13f | Describe any sensitivity analyses conducted to assess robustness of the synthesized results. | 5 |
| Reporting bias assessment | 14 | Describe any methods used to assess risk of bias due to missing results in a synthesis (arising from reporting biases). | 5 |
| Certainty assessment | 15 | Describe any methods used to assess certainty (or confidence) in the body of evidence for an outcome. | 5 |
| **RESULTS** | | |  |
| Study selection | 16a | Describe the results of the search and selection process, from the number of records identified in the search to the number of studies included in the review, ideally using a flow diagram. | Figure 1A |
|  | 16b | Cite studies that might appear to meet the inclusion criteria, but which were excluded, and explain why they were excluded. | Data Extraction Table |
| Study characteristics | 17 | Cite each included study and present its characteristics. | Data Extraction Table |
| Risk of bias in studies | 18 | Present assessments of risk of bias for each included study. | 5-6; Figure 1B |
| Results of individual studies | 19 | For all outcomes, present, for each study: (a) summary statistics for each group (where appropriate) and (b) an effect estimate and its precision (e.g. confidence/credible interval), ideally using structured tables or plots. | Figure 2,3,5; Figure S3-15 |
| Results of syntheses | 20a | For each synthesis, briefly summarise the characteristics and risk of bias among contributing studies. | Figure 2-5; Figure S3-15 |
|  | 20b | Present results of all statistical syntheses conducted. If meta-analysis was done, present for each the summary estimate and its precision (e.g. confidence/credible interval) and measures of statistical heterogeneity. If comparing groups, describe the direction of the effect. | Figure 2-5; Figure S3-15 |
|  | 20c | Present results of all investigations of possible causes of heterogeneity among study results. | 6-8;Figure 2-5; Figure S3-15 |
|  | 20d | Present results of all sensitivity analyses conducted to assess the robustness of the synthesized results. | 8 |
| Reporting biases | 21 | Present assessments of risk of bias due to missing results (arising from reporting biases) for each synthesis assessed. | 8; Figure 4 |
| Certainty of evidence | 22 | Present assessments of certainty (or confidence) in the body of evidence for each outcome assessed. | Figure 2-5; Figure S3-15 |
| **DISCUSSION** | | |  |
| Discussion | 23a | Provide a general interpretation of the results in the context of other evidence. | 10 |
|  | 23b | Discuss any limitations of the evidence included in the review. | 11-12 |
|  | 23c | Discuss any limitations of the review processes used. | 12 |
|  | 23d | Discuss implications of the results for practice, policy, and future research. | 13 |
| **OTHER INFORMATION** | | |  |
| Registration and protocol | 24a | Provide registration information for the review, including register name and registration number, or state that the review was not registered. | 1;4 |
|  | 24b | Indicate where the review protocol can be accessed, or state that a protocol was not prepared. | 4 |
|  | 24c | Describe and explain any amendments to information provided at registration or in the protocol. | 6 |
| Support | 25 | Describe sources of financial or non-financial support for the review, and the role of the funders or sponsors in the review. | 13 |
| Competing interests | 26 | Declare any competing interests of review authors. | 13 |
| Availability of data, code and other materials | 27 | Report which of the following are publicly available and where they can be found: template data collection forms; data extracted from included studies; data used for all analyses; analytic code; any other materials used in the review. | 13 |

*From:*  Page MJ, McKenzie JE, Bossuyt PM, Boutron I, Hoffmann TC, Mulrow CD, et al. The PRISMA 2020 statement: an updated guideline for reporting systematic reviews. BMJ 2021;372:n71. doi: 10.1136/bmj.n

**Supplementary Appendix 2. Database search strategy and number of records returned.**

| Database | Time | Search term | Filter | Number of records identified |
| --- | --- | --- | --- | --- |
| PubMed | Inception to June 4, 2022 | ("cannabi*"[Title/Abstract] OR "marijuana"[Title/Abstract] OR "marihuana"[Title/Abstract] OR "THC"[Title/Abstract] OR "CBD"[Title/Abstract] OR "Tetrahydrocannabinol"[Title/Abstract] OR "win"[Title/Abstract] OR "win55212 2"[Title/Abstract] OR "cp"[Title/Abstract] OR "cp55 940"[Title/Abstract]) AND ("animal*"[Title/Abstract] OR "mouse"[Title/Abstract] OR "mice"[Title/Abstract] OR "rat"[Title/Abstract] OR "rats"[Title/Abstract] OR "rodent*"[Title/Abstract]) AND ("adolescen*"[Title/Abstract] OR “puber*” [Title/Abstract] OR “pubescen*”[Title/Abstract] OR “Juvenil*”[Title/Abstract]) | None | 654 |
| Embase, MEDLINE, APA PsycInfo (Accessed through Ovid) | 1974-Jun 4,2022  1946- June 1, 2022**,**  1806-Jun Week 1, 2022 | ((cannabi* or mari#uana or THC or CBD or tetrahydrocannabinol) and (animal* or mouse or mice or rat or rats or rodent*) and (adolescen* or puber* or pubescen* or juvenil*)).tw. | English language only | 1687 |

**Supplementary Appendix 3. Further details for methods.**

**Study inclusion criteria according to the PICO method**

1. **Study characteristics**: Original research studies published in peer-reviewed journals (excluding reviews, opinions, conference abstracts, theses, etc), written in English, published from inception to the date of search, and full-text available online.

2. **Animals**: Mice or rats of any strain, either wildtype or genetically modified, either or both sexes.

3. **Intervention**:

1) ***Substance***: THC and synthetic cannabinoid type 1 receptor (CB1R) agonists (e.g., WIN 55, 212-2) as well as CBD.

2) ***Drug treatment***: Chronic administration (repeated exposure for more than three days) at any dose, through any route (e.g., subcutaneous, intraperitoneal, inhaled, oral), either at random or at regular frequency; Onset of drug treatment between postnatal day (PND) 21 and PND56. We excluded experiments that combined cannabinoids with other interventions (e.g., alcohol co-treatment, maternal deprivation) without a cannabinoid-only group. Studies with genetic modifications addressing gene-environment interaction hypotheses were treated separately.

3) ***Control group treatment:*** Age-matched conspecifics of the same genotype, treated with vehicle or placebo through the same route and administration paradigm as the experimental group(s).

4). ***Outcome measures*:** With a focus on the protracted effects of cannabinoid exposure, we included behavioural analyses performed shortly after chronic treatment cessation (short-term - 24 hours to 10 days after the final dose) or after long-term abstinence (at least 11 days after treatment). Studies that solely examined the acute effects (within 24 hours after exposure) were not included in our analysis.

To assess the effects of cannabinoid exposure on schizophrenia-related phenotypes, we included 11 common rodent behavioural tests (Table 1) which cover the three core domains of schizophrenia symptoms: positive symptoms (such as hallucinations and delusions), negative symptoms (such as social withdrawal and anhedonia), and cognitive impairments (such as memory dysfunction). To standardize the outcome measures, we designated a primary behavioural readout for each behavioural test, as listed in Table 1. If the specified outcome measure was not reported from an experiment, we evaluated on a case-by-case basis whether an alternative measure is feasible, and all alternative outcomes measures included were recorded. Studies without appropriate outcome measures were excluded from analysis.

The following formula were used to calculate the relevant outcome measures.

$$Discrimination index \left( DI \right)=\frac{\left[ time exploring novel object /location- time exploring familiar object/location \right]}{\left[ total time exploring both novel and familiar objects \right]}$$

$$Social preference index=\frac{\left[ time exploring novel conspecific- time exploring familiar conspecific \right]}{\left[ total time exploring both novel and familiar conspecific \right]}$$

$$Social motivation index=\frac{\left[ time exploring novel conspecific- time exploring novel object \right]}{\left[ total time exploring both conspecific and object \right]}$$

$$Sucrose preference index=\frac{\left[ sucrose solution intake(g) \right]}{\left[ sucrose solution consumption (g)+ water consumption(g) \right]}\times100 \%$$

$$\%Correct alternations =\frac{\left[ number of alternations \right]}{\left[ total number of arm entries - 2 \right]}\times100\%$$

$$PPI \left( \% \right)=\frac{\left[ 1- average startle amplitude to pulse with prepulse \right]}{\left[ average startle amplitude to pulse only \right]}\times100\%$$

**Data Extraction strategy**

From each experiment, we extracted sample sizes per group, mean values of outcomes and standard errors/distributions. Alternatively, when the mean was not reported, we extracted sample sizes per group, and t statistics. To avoid double-counting, if a single control group was used for multiple treatment groups, the sample size of the control group was divided by the number of experiment groups it serves. When essential numerical data were not available from texts, we used a digital ruler software (WebPlotDigitizer Ver. 4.6, Ankit Rohatgi) to extract data from figures and diagrams. Furthermore, if key information could not be obtained from the articles, we attempted to contact authors by email (max. 2 attempts). Sources of data collected from each experiment were documented. In cases where a behavioural outcome was repeatedly measured from the same cohort of animals at different times, data from the time point with the largest effect of drug treatment was included. Whenever the articles gave a sampling range rather than the precise number of animals in a group, we applied the lowest possible value within the specified range. To calculate mean discrimination indexes and standard errors from studies reporting only mean time and standard errors per group, we performed a bootstrapping procedure with 10,000 iterations to create simulated individual discrimination index samples. Any discrepancies in the data extracted were further checked and discussed, and additional arbitrators (DM, MDF) adjudicated unresolved disagreements.

**Supplementary Appendix 4**. **Descriptive summary studies that assessed gene-environment interactions**

Human epidemiological studies have established a clear link between cannabis use and psychosis. However, not all cannabis users develop psychosis, suggesting that genetic factors may influence the risk of psychosis in response to cannabis exposure. Although the gene and environmental interaction of schizophrenia has been studied in mouse models to some detail, studies are still scarce in the context of cannabis exposure and genetic vulnerability. In this systematic review, we identified only a handful of studies that investigated the gene x environment interactions in schizophrenia by exposing adolescent genetically modified mice to CB1R agonists. These studies were not included in the main analysis, a descriptive summary of the studies is presented here.

Table of the existing studies approaching adolescent cannabinoid exposure and schizophrenia in a mouse gene-environment interaction model:

| Study | Species | Genotype | Associated pathway | Sex | Administration | Age of exposure | Behavioural task | Behavioural outcome in GxE model | Significant GxE interaction |
| --- | --- | --- | --- | --- | --- | --- | --- | --- | --- |
| [Jouroukhin et al.(2019)](https://www.biologicalpsychiatryjournal.com/article/S0006-3223(18)31746-3/fulltext) | Mouse | astrocyte DN-DISC1 -/- | NF-κB-COX-2 signalling | M | THC | P30-51 | Y maze | impaired spatial working memory | yes |
|  |  |  |  |  |  |  | Novel object | impaired object recognition memory | yes |
|  |  |  |  |  |  |  | Novel object location | impaired object location memory | no |
|  |  |  |  | F |  |  | Y maze | impaired spatial working memory | yes |
|  |  |  |  |  |  |  | Novel object | no significant effect | no |
|  |  |  |  |  |  |  | Novel object location | no significant effect | no |
| [Segal-Gavish et al.(2017)](https://academic.oup.com/hmg/article/26/13/2462/3574683) | Mouse | forebrain pyramidal neuron DN-DISC1 -/- | BDNF | M | THC | P42-51 | Open Field | no significant effect on locomotion | no |
|  |  |  |  |  |  |  | Novel object | impaired object recognition memory | yes |
| [Ballinger et al.(2015)](https://www.sciencedirect.com/science/article/abs/pii/S0969996115002247?via%3Dihub) | Mouse | forebrain pyramidal neuron DN-DISC1 -/- | CB1R expression in the PFC and AMG | M | THC | P28-48 | Y maze | no significant effect | no |
|  |  |  |  |  |  |  | Novel object | no significant effect | no |
|  |  |  |  |  |  |  | Fear Conditioning | impaired contextual memory | no |
| [Long et al.(2013)](https://academic.oup.com/ijnp/article/16/1/163/628820) | Mouse | Nrg1 +/- | CB1R and 5-HT2AR binding | M | THC | ~P31-50 | Open Field | lower locomotion | no |
|  |  |  |  | M | THC |  | Prepulse Inhibition | no significant effect on %PPI | no |
| O’Tuathaigh et al.(2012) | Mouse | COMT -/- | catabolism of cortical dopamine | M | WIN55,212–2  2.5mg/kg | P32-52 | Prepulse Inhibition | no significant effect on %PPI | no |
|  |  |  |  |  |  |  | Social preference | no significant effect | no |
|  |  |  |  |  |  |  | Social motivation | no significant effect | yes |
| [O’Tuathaigh et al.(2010)](https://www.nature.com/articles/npp2010100) | Mouse | COMT -/- | catabolism of cortical dopamine | M | THC | P32-52 | Open Field | increased locomotion | no |
|  |  |  |  |  |  |  | Y maze | impaired spatial working memory | yes |
|  |  |  |  |  |  |  | Novel object | impaired object recognition memory | no |
|  |  |  |  |  |  |  | Social motivation | no significant effect | no |
|  |  |  |  |  |  |  | Social preference | decreased social novelty preference | no |
|  |  |  |  | F |  |  | Open Field | no significant effect | no |
|  |  |  |  |  |  |  | Y maze | impaired spatial working memory | no |
|  |  |  |  |  |  |  | Novel object | impaired object recognition memory | no |
|  |  |  |  |  |  |  | Social motivation | no significant effect | no |
|  |  |  |  |  |  |  | Social preference | decreased social novelty preference | no |

**Supplementary Appendix 5.** **Heterogeneity with and without outliers for CB1R agonists and CBD**.

|  | **OFT** | | **WM** | | **NOR** | | **NOL** | | **SNP** | | **SM** | | **PPI** | | **SPT** | | **FC** | |
| --- | --- | --- | --- | --- | --- | --- | --- | --- | --- | --- | --- | --- | --- | --- | --- | --- | --- | --- |
|  | Agonists | CBD | Agonists | CBD | Agonists | CBD | Agonists | CBD | Agonists | CBD | Agonists | CBD | Agonists | CBD | Agonists | CBD | Agonists | CBD |
| ES | -0.11 | -0.23 | -0.58 | -0.55 | -0.63 | -0.44 | -0.7 | -0.47 | -0.92 | - | -0.4 | -0.71 | -0.48 | 0.4 | -0.92 | 0.1 | -0.45 | 0.53 |
| *I*^2^_main_ | 54% | 0% | 60% | - | 71% | 9% | 69% | - | 71% | - | 0% | - | 76% | 48% | 70% | 21% | 46% | 0% |
| 95% CI | 40-64% | 0-58% | 38-75% | - | 63-78% | 0-86% | 53-80% | - | 53-82% | - | 0-58% | - | 66-84% | 0-80% | 39-85% | 0-66% | 0-73% | 0-68% |
| N_total_ | 75 | 12 | 24 | 1 | 56 | 4 | 22 | 1 | 17 | 0 | 12 | 1 | 27 | 6 | 9 | 5 | 11 | 8 |
| N _outliers_ | 12 | 0 | 1 | 0 | 9 | 0 | 4 | 0 | 1 | - | 0 | - | 5 | 0 | 1 | 0 | 0 | 0 |
| ES_no outlier_ | -0.153 | - | -0.653 | - | -0.435 | - | -0.7087 | - | -0.677 | - | - | - | -0.477 | - | -0.58 | - | - | - |
| p | 0.0114 | - | 0.0009 | - | < 0.0001 | - | < 0.0001 | - | 0.0086 | - | - | - | 0.0069 | - | 0.0573 | - | - | - |
| *I*^2^_no outliers_ | 0% | - | 50% | - | 47% | - | 26% | - | 66% | - | - | - | 60% | - | 44% | - | - | - |
| 95% CI | 0-29% | - | 18-69% | - | 26-62% | - | 0-59% | - | 42-80% | - | - | - | 36-75% | - | 0-75% | - | - | - |

Table shows the effect sizes for main analysis (ES), heterogeneity (I^2^_main_ and corresponding 95%CI), total number of data points for each behaviour test (N_total_), number of outliers (N_outliers_), effect sizes after outliers are removed (ES_no outliers_) and associated statistical significance (p), heterogeneity after outliers are removed (I^2^_no outliers_ and corresponding 95%CI). A data point was regarded as an outlier if its 95% conﬁdence interval does not overlap with that of the pooled effect. OF-novelty induced hyperactivity; WM- working memory; NOR- novel object recognition; NOL- novel object location; SNP- social novelty preference; PPI- prepulse inhibition, FC- fear conditioning; SM- social motivation; SP- sucrose preference.

**Figure S1 Subgroup analysis on dose of THC administered.**

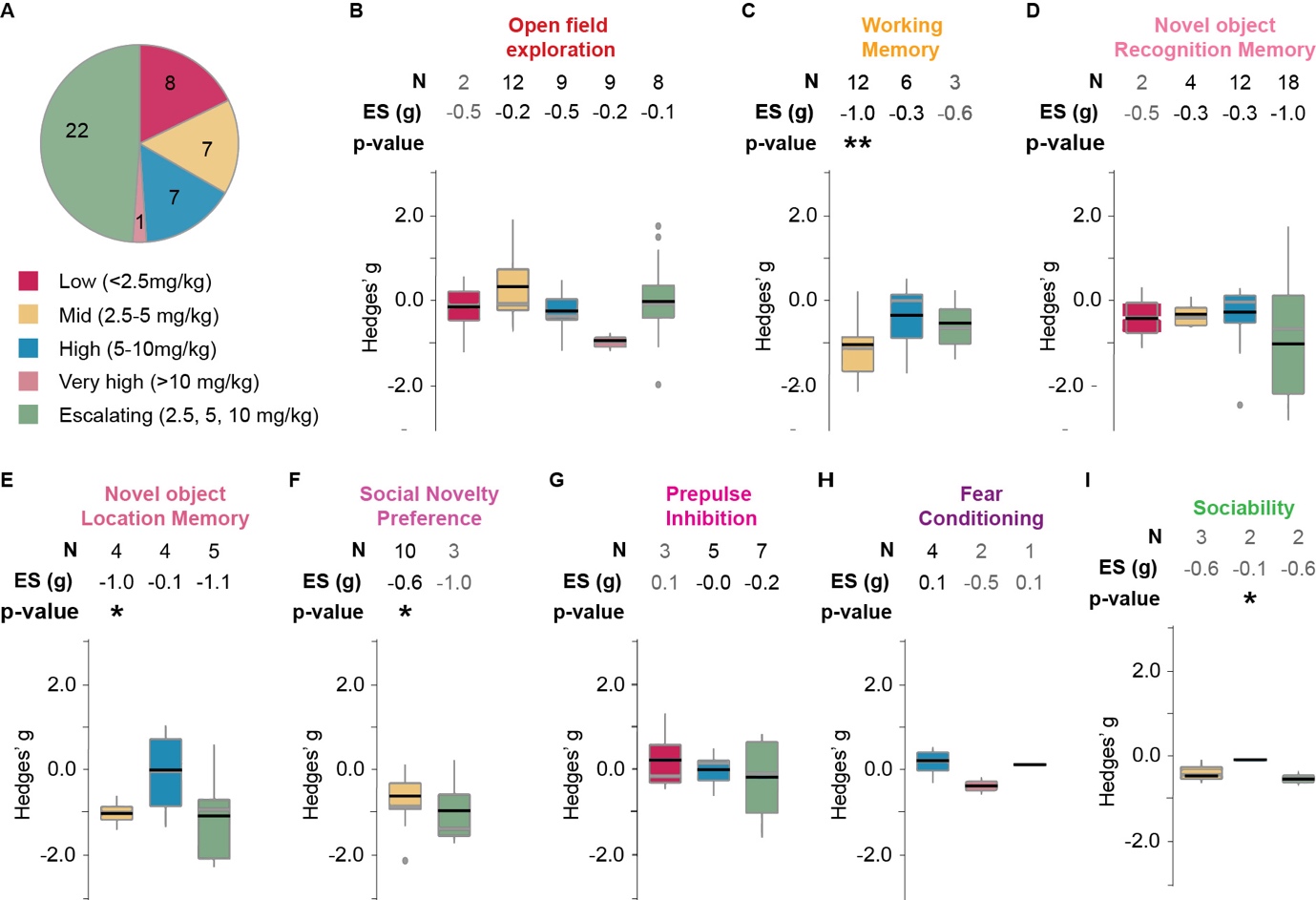

**A.** Proportion of different doses used in the studies included in the quantitative analysis. **B-I**. Boxplots displaying the distribution of effect sizes (Hedge’s g) for THC in each behavioural test sub-grouped by dose. Number of experiments included (N), pooled effect size (ES) and corresponding significance levels are listed on top of each corresponding boxplot. * p<0.05, **p<0.005, *** p<0.0005. Black horizontal lines in the boxplots indicate pooled subgroup effect sizes (inverse variance weighted, mixed effect model), grey horizontal lines indicate medians. Outliers are indicated by grey dots. OF- locomotion in a novel open field; WM- working memory; NOR- novel object recognition; NOL- novel object location; SNP- social novelty preference; PPI- pre-pulse inhibition, FC- fear conditioning; SM- social motivation; SP- sucrose preference.

**Figure S2 Subgroup analysis on timing of administration onset of THC**
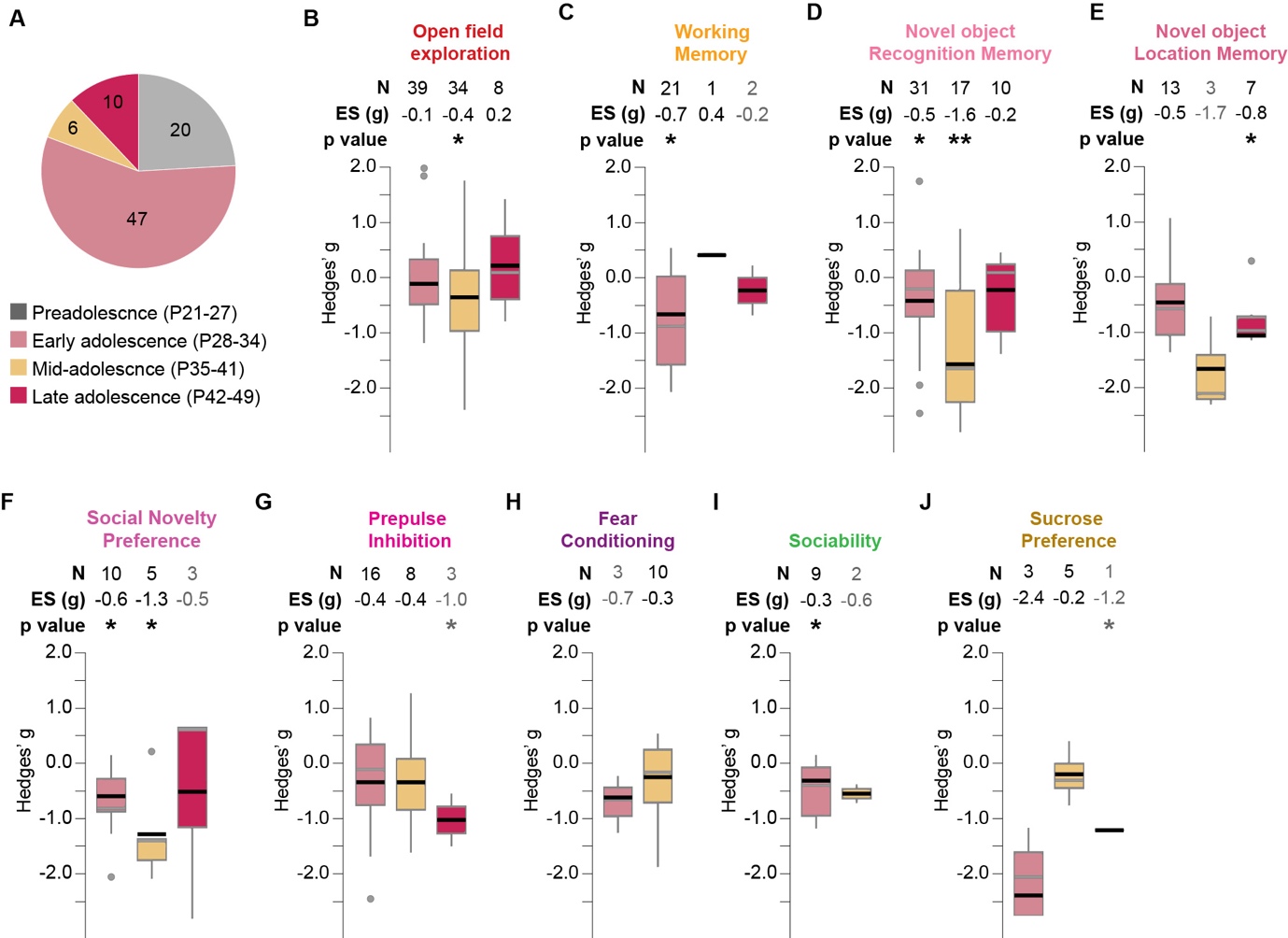

**A.** Proportion of different administration onset used in the studies included in the quantitative analysis. **B-I**. Boxplots displaying the distribution of effect sizes (Hedge’s g) for THC in each behavioural test sub-grouped by onset of treatment. Number of experiments included (N), pooled effect size (ES) and corresponding significance levels are listed on top of each corresponding boxplot. * p<0.05, **p<0.005, *** p<0.0005. Black horizontal lines in the boxplots indicate pooled subgroup effect sizes (inverse variance weighted, mixed effect model), grey horizontal lines indicate medians. Outliers are indicated by grey dots. OF- locomotion in a novel open field; WM- working memory; NOR- novel object recognition; NOL- novel object location; SNP- social novelty preference; PPI- pre-pulse inhibition, FC- fear conditioning; SM- social motivation; SP- sucrose preference.

**Figure S3. Effect of CB1R agonists on locomotion in the open field test.**

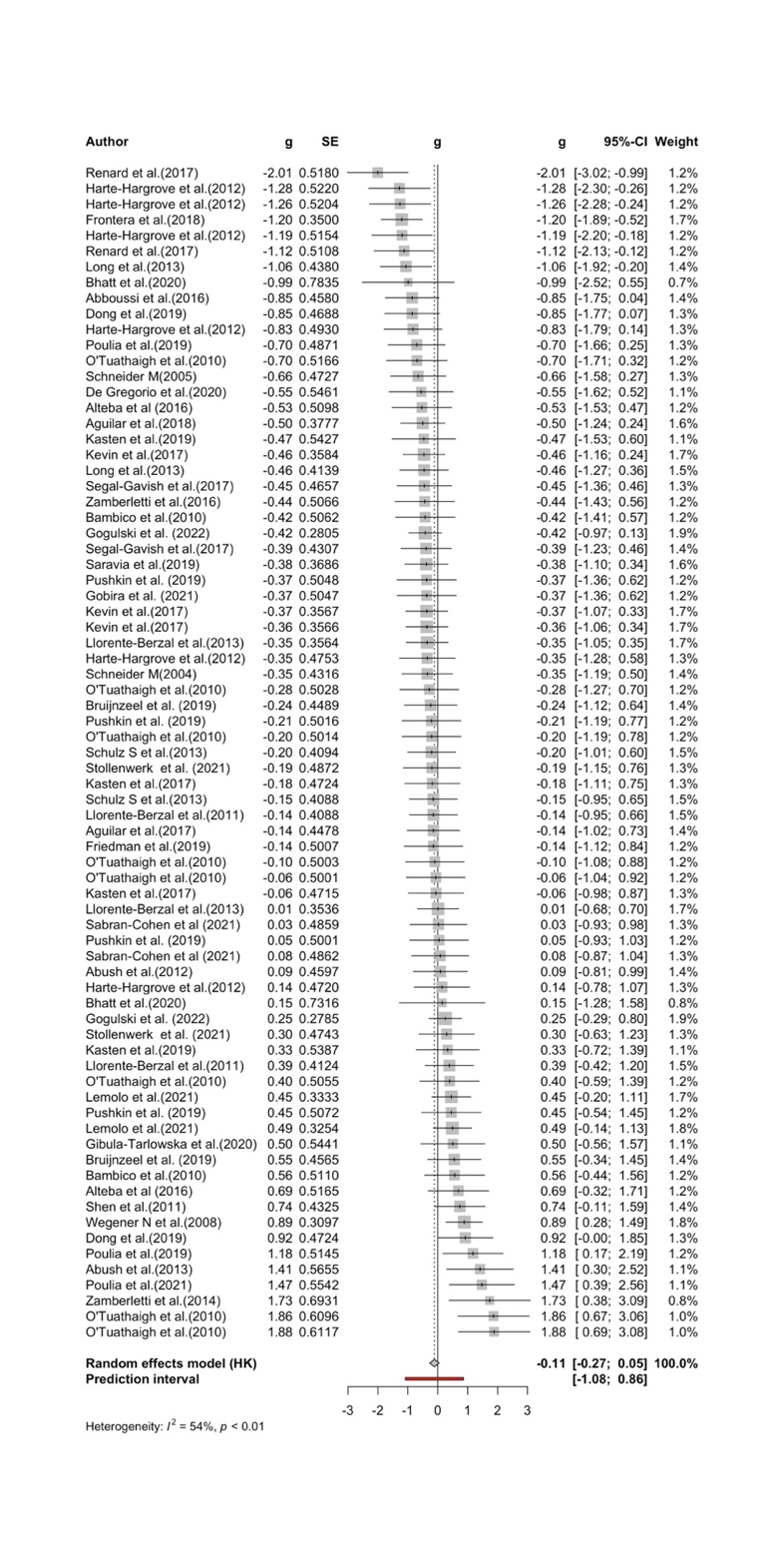

**Forest plot** showing the standardised mean difference (g) in locomotion between the CB1R agonist treatment and control groups for each study, along with the 95% confidence intervals (95%-CIs). The size of the squares represents the weight of each study in the meta-analysis. The horizontal lines across the squares represent 95%-Cis of individual effects. The diamond at the bottom represents the pooled effect size and 95%-CI across all studies. The red horizontal line indicates the prediction interval.

**Figure S4. Effect of CB1R agonists on working memory**

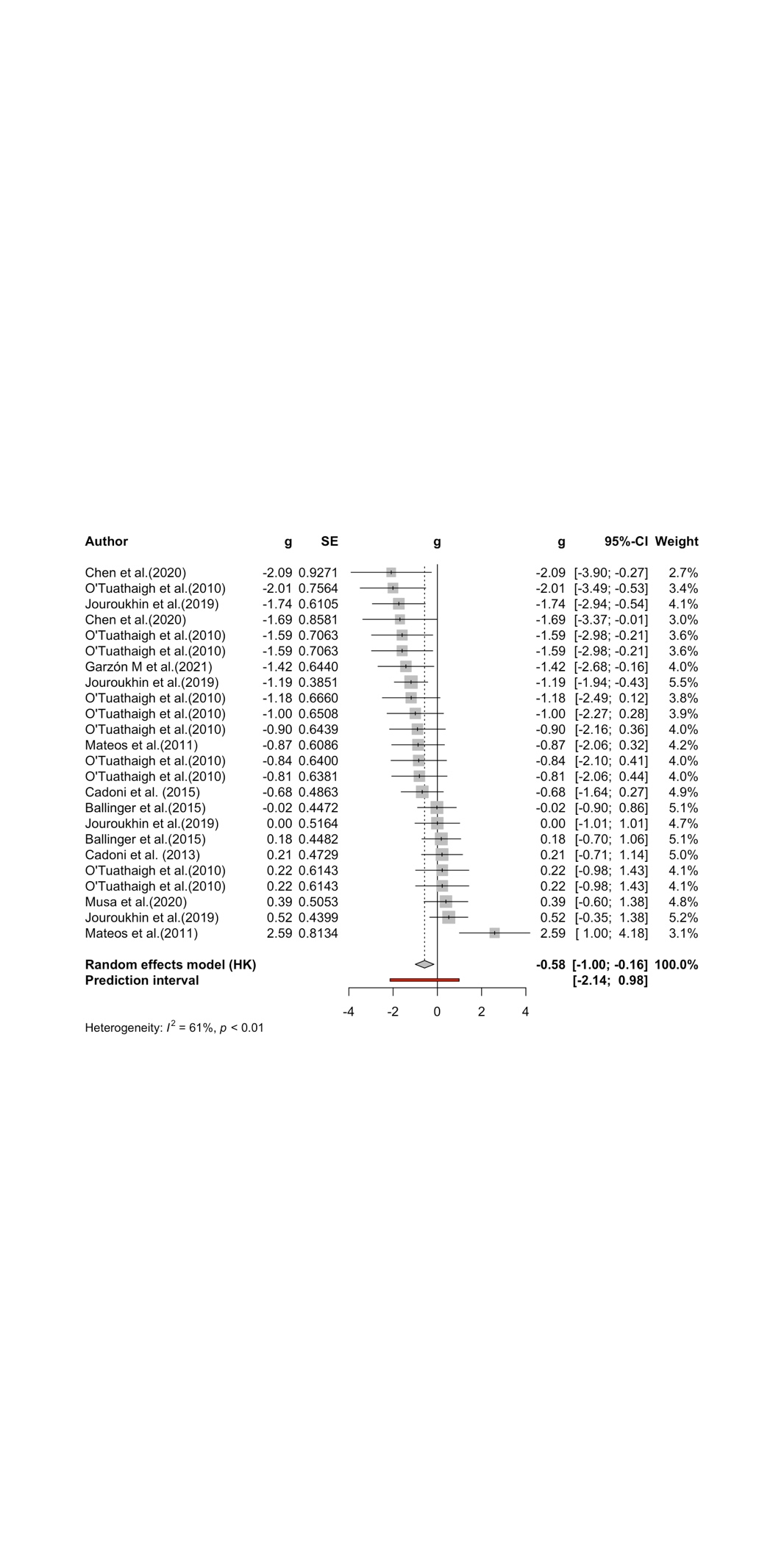

**Forest plot** showing the standardised mean difference (g) in %correct alternation between the CB1R agonist treatment and control groups for each study, along with the 95% confidence intervals (95%-CIs). The size of the squares represents the weight of each study in the meta-analysis. The horizontal lines across the squares represent 95%-Cis of individual effects. The diamond at the bottom represents the pooled effect size and 95%-CI across all studies. The red horizontal line indicates the prediction interval.

**Figure S5.** **Effect of CB1R agonists on** **discrimination index in the novel object recognition test**

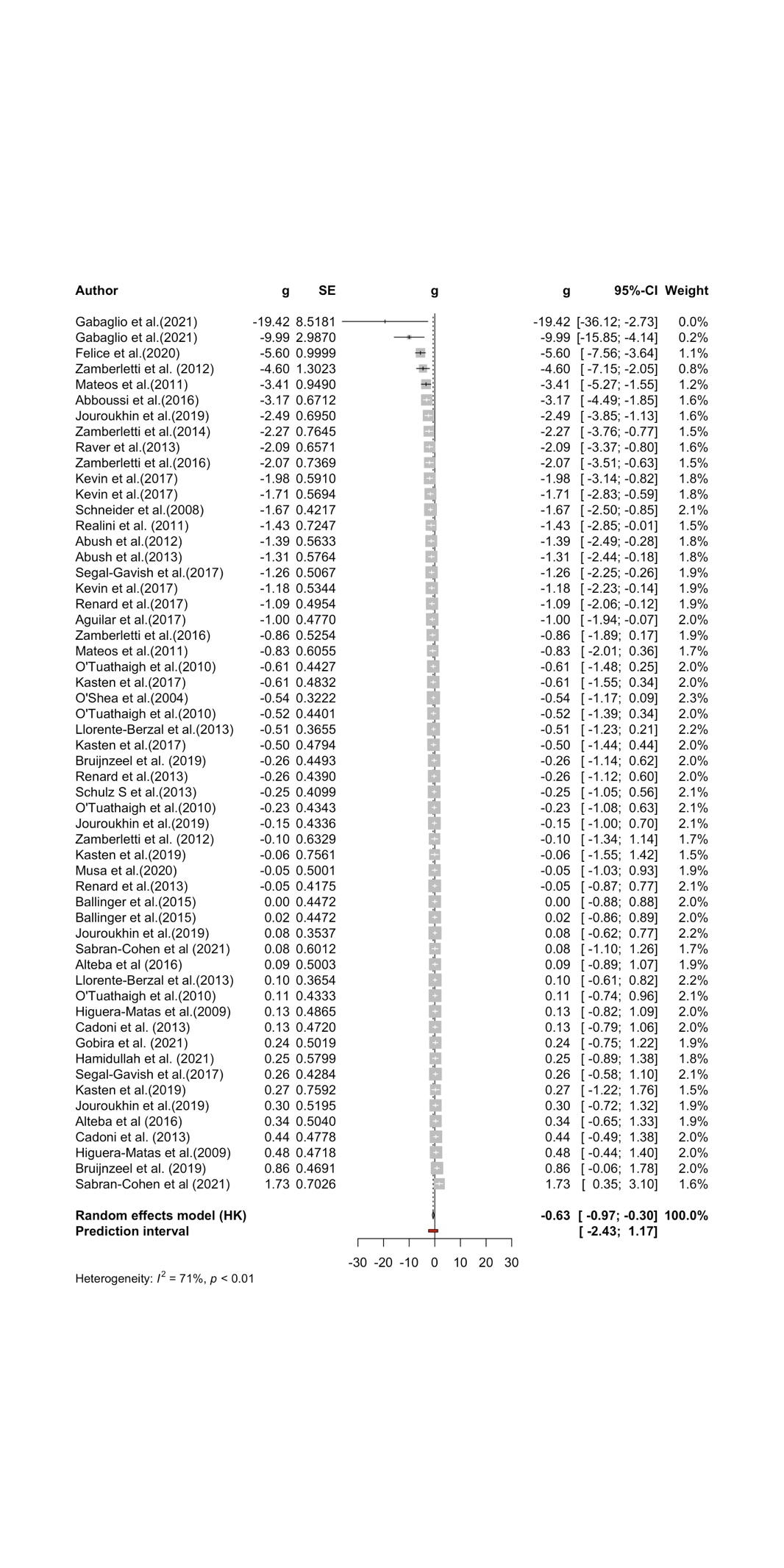

**Forest plot** showing the standardised mean difference (g) in discrimination index between the CB1R agonist treatment and control groups for each study, along with the 95% confidence intervals (95%-CIs). The size of the squares represents the weight of each study in the meta-analysis. The horizontal lines across the squares represent 95%-Cis of individual effects. The diamond at the bottom represents the pooled effect size and 95%-CI across all studies. The red horizontal line indicates the prediction interval.

**Figure S6.** **Effect of CB1R agonists on discrimination index in the novel object location test**

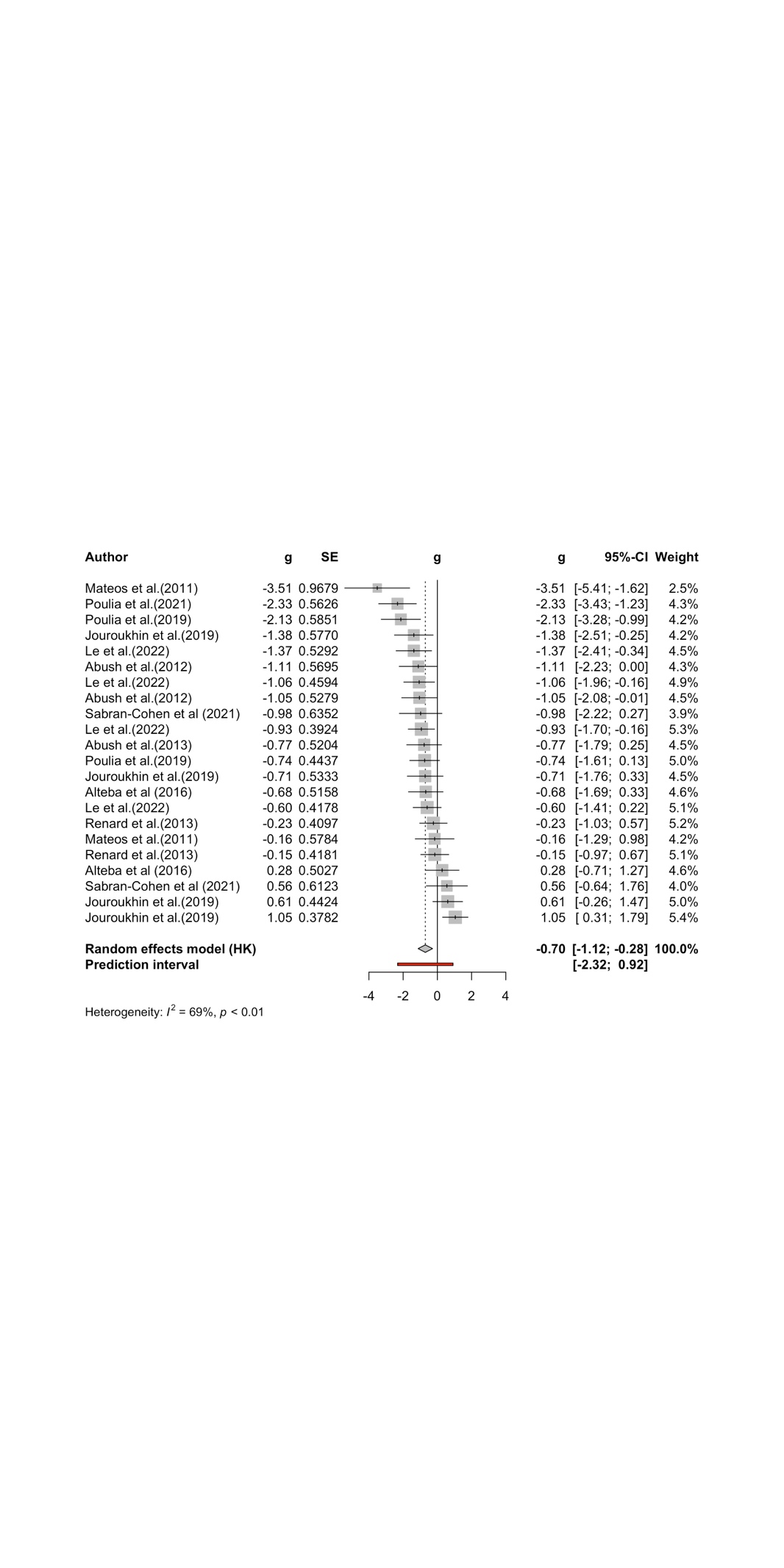

**Forest plot** showing the standardised mean difference (g) in discrimination index between the CB1R agonist treatment and control groups for each study, along with the 95% confidence intervals (95%-CIs). The size of the squares represents the weight of each study in the meta-analysis. The horizontal lines across the squares represent 95%-Cis of individual effects. The diamond at the bottom represents the pooled effect size and 95%-CI across all studies. The red horizontal line indicates the prediction interval.

**Figure S7.** **Effect of CB1R agonists in the social novelty preference test**

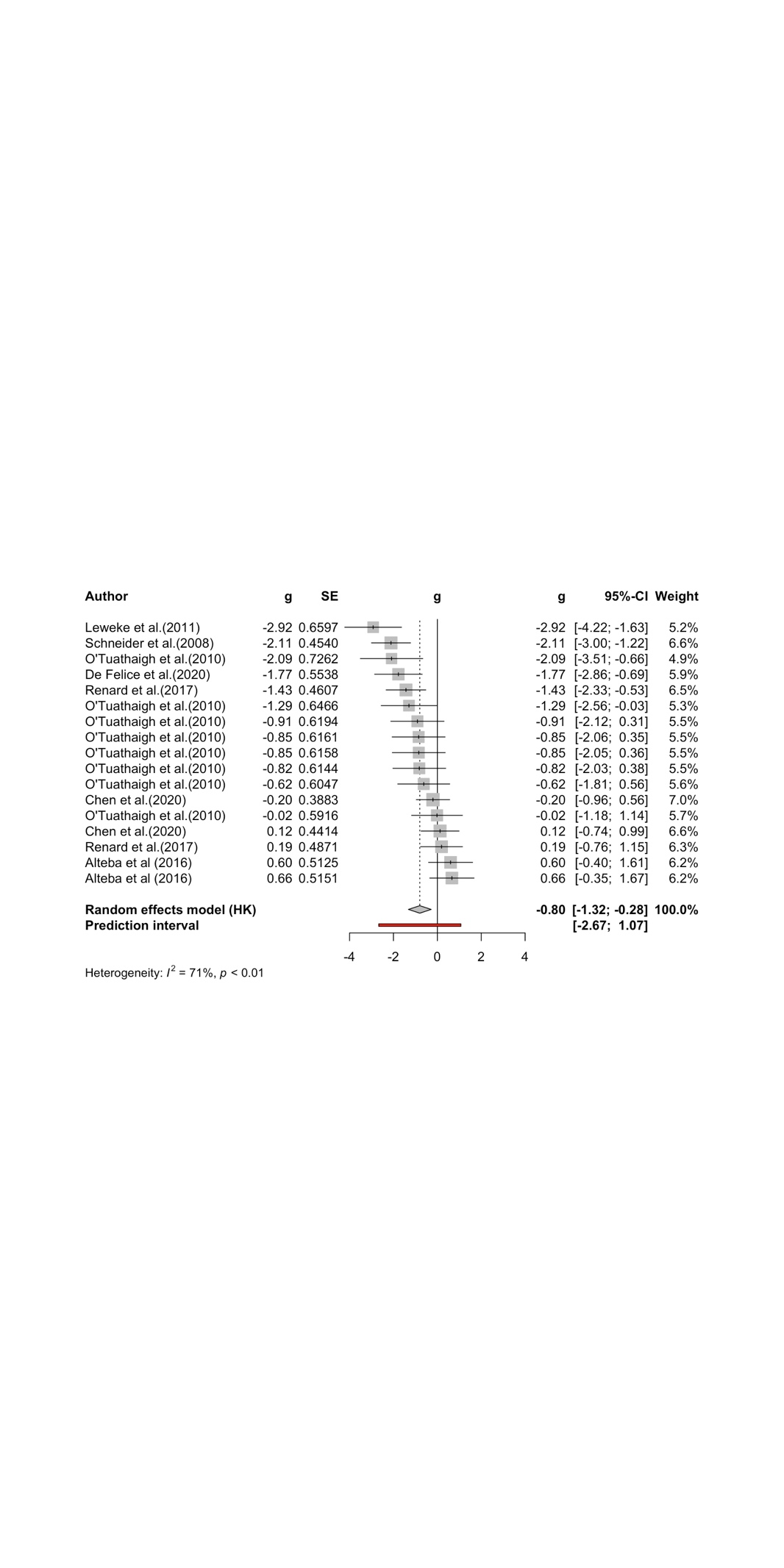

**Forest plot** showing the standardised mean difference (g) in social preference index between the CB1R agonist treatment and control groups for each study, along with the 95% confidence intervals (95%-CIs). The size of the squares represents the weight of each study in the meta-analysis. The horizontal lines across the squares represent 95%-Cis of individual effects. The diamond at the bottom represents the pooled effect size and 95%-CI across all studies. The red horizontal line indicates the prediction interval.

**Figure S8. Effect of CB1R agonists on pre-pulse inhibition**

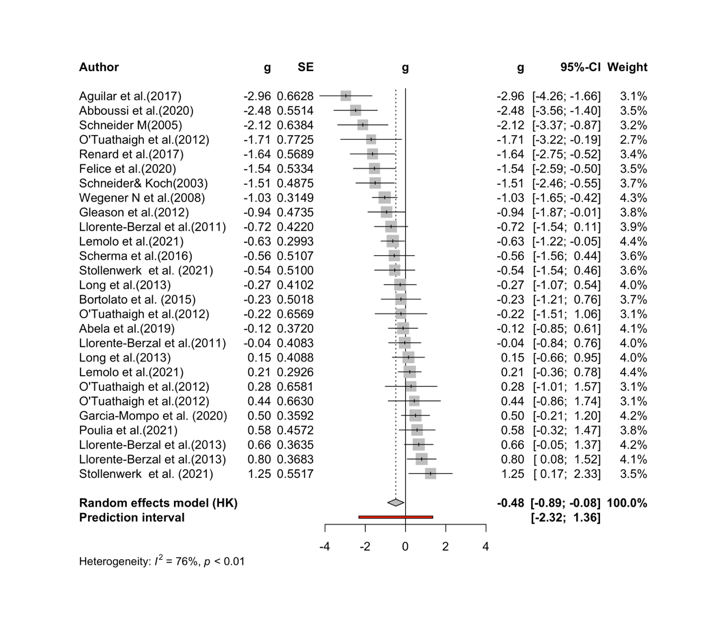

**Forest plot** showing the standardised mean difference (g) in %PPI between the CB1R agonist treatment and control groups for each study, along with the 95% confidence intervals (95%-CIs). The size of the squares represents the weight of each study in the meta-analysis. The horizontal lines across the squares represent 95%-Cis of individual effects. The diamond at the bottom represents the pooled effect size and 95%-CI across all studies. The red horizontal line indicates the prediction interval.

**Figure S9. Effect of CB1R agonists on** **fear memory recall**

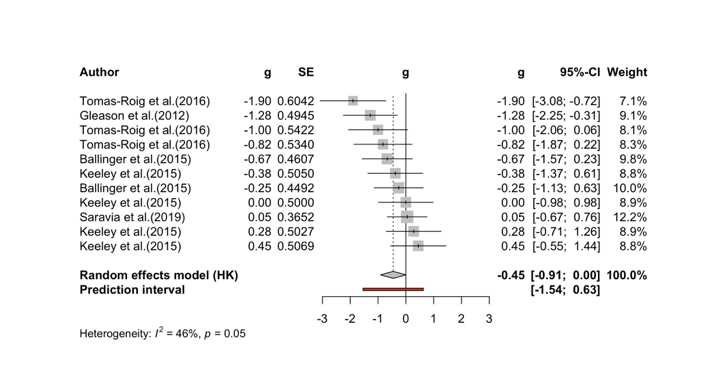

**Forest plot** showing the standardised mean difference (g) in %time freezing in recall trial between the CB1R agonist treatment and control groups for each study, along with the 95% confidence intervals (95%-CIs). The size of the squares represents the weight of each study in the meta-analysis. The horizontal lines across the squares represent 95%-Cis of individual effects. The diamond at the bottom represents the pooled effect size and 95%-CI across all studies. The red horizontal line indicates the prediction interval.

**Figure S10. Effect of CB1R agonists on social motivation**

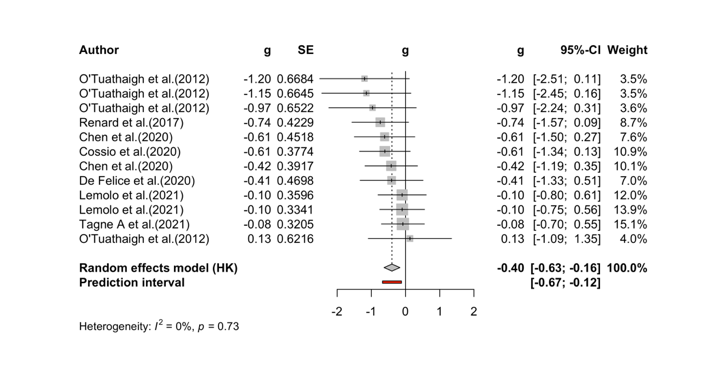

**Forest plot** showing the standardised mean difference (g) in social motivation index in recall trial between the CB1R agonist treatment and control groups for each study, along with the 95% confidence intervals (95%-CIs). The size of the squares represents the weight of each study in the meta-analysis. The horizontal lines across the squares represent 95%-Cis of individual effects. The diamond at the bottom represents the pooled effect size and 95%-CI across all studies. The red horizontal line indicates the prediction interval.

**Figure S11. Effect of CB1R agonists on sucrose preference**

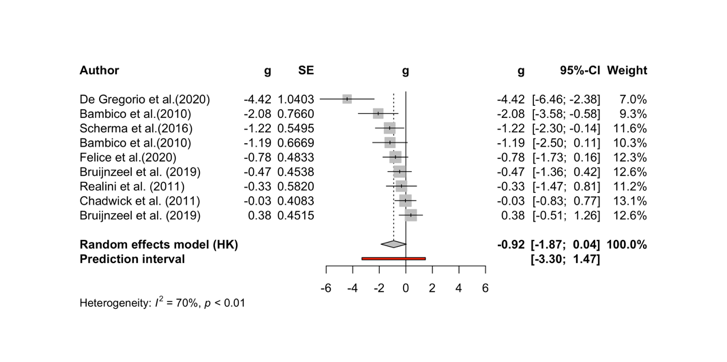

**Forest plot** showing the standardised mean difference (g) in sucrose preference index in recall trial between the CB1R agonist treatment and control groups for each study, along with the 95% confidence intervals (95%-CIs). The size of the squares represents the weight of each study in the meta-analysis. The horizontal lines across the squares represent 95%-Cis of individual effects. The diamond at the bottom represents the pooled effect size and 95%-CI across all studies. The red horizontal line indicates the prediction interval.

**Figure S12. Effect of CBD on locomotion in the open field test**

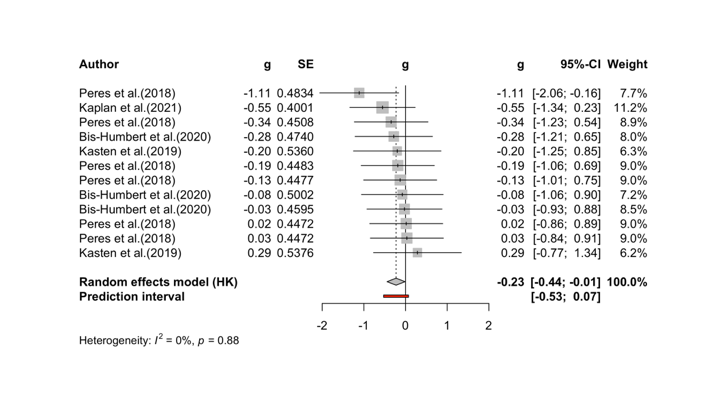

**Forest plot** showing the standardised mean difference (g) in locomotion between the CBD treatment and control groups for each study, along with the 95% confidence intervals (95%-CIs). The size of the squares represents the weight of each study in the meta-analysis. The horizontal lines across the squares represent 95%-Cis of individual effects. The diamond at the bottom represents the pooled effect size and 95%-CI across all studies. The red horizontal line indicates the prediction interval.

**Figure S13. Effect of CBD on discrimination index in the novel object recognition test**

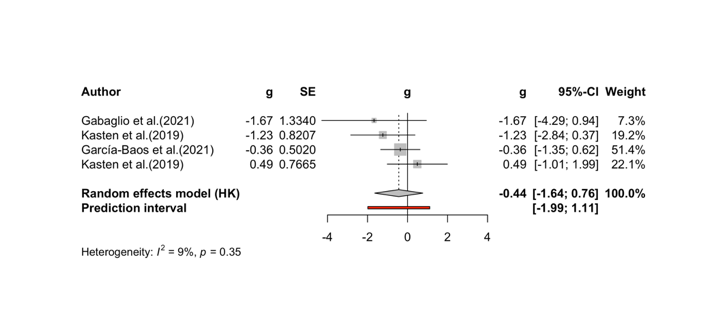

**Forest plot** showing the standardised mean difference (g) in discrimination index between the CBD treatment and control groups for each study, along with the 95% confidence intervals (95%-CIs). The size of the squares represents the weight of each study in the meta-analysis. The horizontal lines across the squares represent 95%-Cis of individual effects. The diamond at the bottom represents the pooled effect size and 95%-CI across all studies. The red horizontal line indicates the prediction interval.

**Figure S14. Effect of CBD on pre-pulse inhibition**

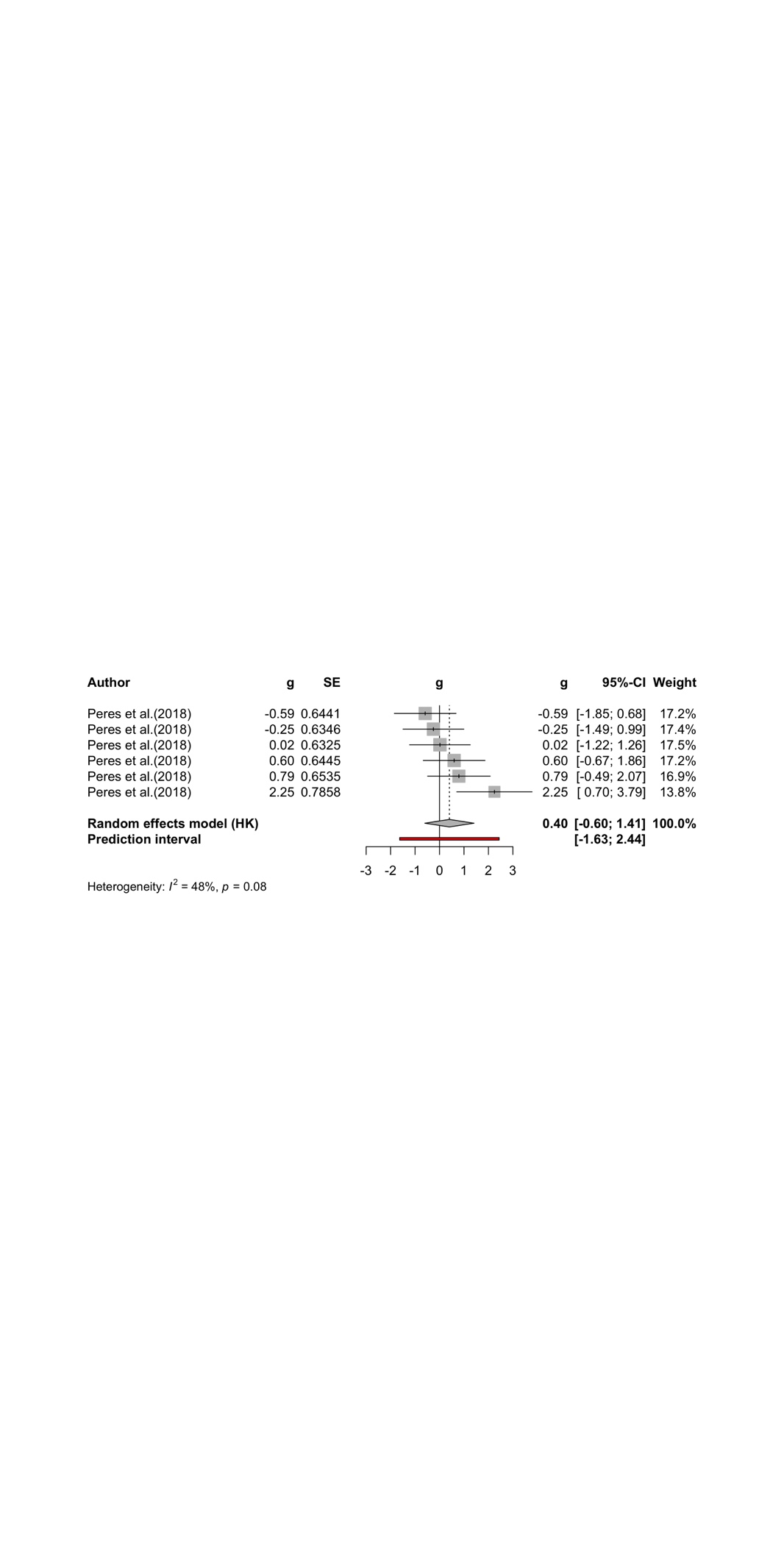

**Forest plot** showing the standardised mean difference (g) in %PPI between the CBD treatment and control groups for each study, along with the 95% confidence intervals (95%-CIs). The size of the squares represents the weight of each study in the meta-analysis. The horizontal lines across the squares represent 95%-Cis of individual effects. The diamond at the bottom represents the pooled effect size and 95%-CI across all studies. The red horizontal line indicates the prediction interval.

**Figure S15. Effect of CBD on fear memory recall**

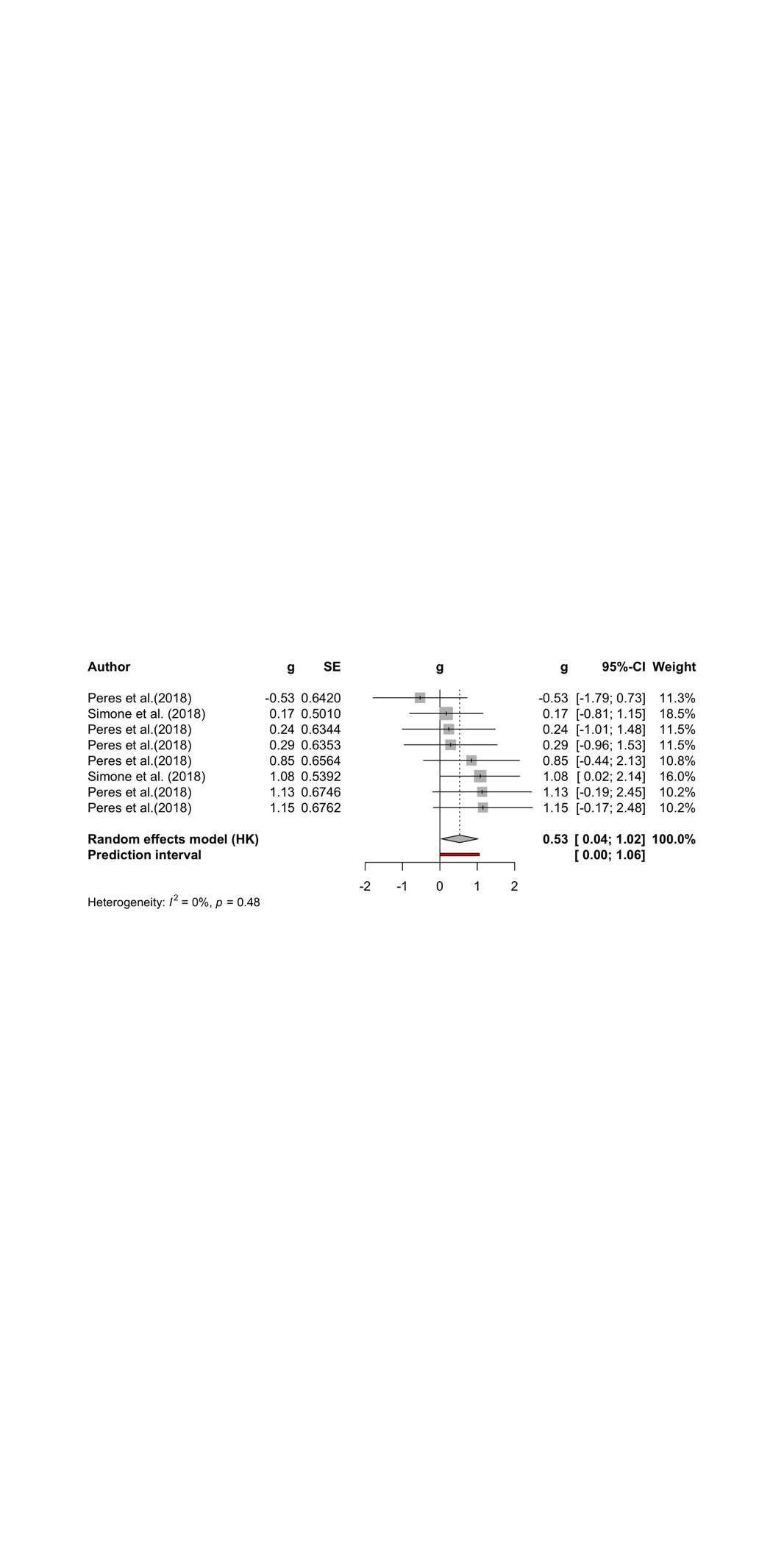

**Forest plot** showing the standardised mean difference (g) in %time freezing in recall trial between the CBD treatment and control groups for each study, along with the 95% confidence intervals (95%-CIs). The size of the squares represents the weight of each study in the meta-analysis. The horizontal lines across the squares represent 95%-Cis of individual effects. The diamond at the bottom represents the pooled effect size and 95%-CI across all studies. The red horizontal line indicates the prediction interval.
